# Photoreceptors have a dual dependency on both aerobic glycolysis and OXPHOS and diverge metabolically from other retinal neurons

**DOI:** 10.64898/2026.03.02.709047

**Authors:** Gabriele M. Wögenstein, Luca Ravotto, Vyara Todorova, Rachel M. Meister, Marijana Samardzija, Bruno Weber, Christian Grimm

## Abstract

The retina metabolizes glucose into lactate, a hallmark of aerobic glycolysis known as the Warburg effect. Although evidence points to rod photoreceptors as the primary source of aerobic glycolysis, a comparison of the energy metabolism in different retinal neurons has yet to be performed. We combined two-photon fluorescence lifetime imaging of biosensors with pharmacological protocols to analyse metabolic dynamics in healthy and diseased photoreceptors and in inner retinal neurons. Our data reveal distinct metabolic profiles among retinal neurons, identify rods as the drivers of aerobic glycolysis, demonstrate that inner retinal neurons rely on oxidative phosphorylation, show that rods need both glycolysis and oxidative phosphorylation to maintain ATP levels, and suggest that rods can metabolize lactate. A mutation causing retinitis pigmentosa increases lactate production in rods but changes the energy metabolism only subtly otherwise. Our results improve the understanding of retinal physiology and are relevant for pathologies involving imbalanced energy metabolism.

## Introduction

The retina converts light into electrochemical signals that are relayed to the brain via the optic nerve. As the required cellular processes are energetically costly, the retina has a very high metabolic demand (*1, 2*). Inner retinal neurons primarily use energy for synaptic transmission, ion pumping and neurotransmitter recycling (*3*). Photoreceptors in the outer retina demand high energy levels to maintain ion homeostasis (the dark current), regulate the phototransduction cascade, sustain glutamate release at synaptic terminals, and support the continuous renewal of their outer segments (*4–7*). To meet its energy demands and generate ATP, the outer retina consumes oxygen and glucose at very high rates (*8, 9*). Although most cells generate ATP primarily through oxidative phosphorylation (OXPHOS) in mitochondria, Otto Warburg discovered a century ago that retinal cells, like cancer cells, ferment glucose to lactate in the presence of oxygen - a process known as aerobic glycolysis or the Warburg effect (*10, 11*). Since then, considerable efforts have been devoted to describing retinal energy metabolism in detail and defining the cellular source of lactate in the retina (*12*). Based on a body of data including the correlation of reduced expression of enzymes involved in aerobic glycolysis, such as hexokinase 2 (HK2), lactate dehydrogenase (LDH) A, and pyruvate kinase M2 (PKM2) with impaired photoreceptor integrity and function (*13–22*), photoreceptors have been suggested to be the main producers of retinal lactate. However, most conclusions are based on genetic models, protein and gene expression data and/or C13 tracing experiments using whole retinas (*13–25*), and only a few studies are available that directly measure metabolites in specific cell types of the retina (*26–29*).

Aerobic glycolysis with its high glucose consumption rate can facilitate the incorporation of nutrients into biomass to meet high anabolic demands (*30*). This process may be preferred by photoreceptors over OXPHOS because it satisfies immediate ATP demands while simultaneously supplying the precursor molecules needed for the maintenance of cell structures, including diurnal outer segment renewal (*13, 31*). The integration of experimental data with the theoretical framework has recently led to a model of the metabolic ecosystem in the retina. This model states that glucose from the choroidal blood is transported through the retinal pigment epithelium (RPE) and delivered to photoreceptors (*32*). There, it is metabolized via aerobic glycolysis, generating ATP and the molecules required for lipid, protein and nucleic acid biosynthesis, as well as lactate, which is the end product of the process (*33, 34*). This lactate is then released into the extracellular space and taken up by the RPE, where it is used to produce energy via OXPHOS (*12, 24, 25, 35*).

Inner retinal signal processing involves three types of interneurons (bipolar, horizontal and amacrine cells) before the signals reach the retinal ganglion cells (RGCs), the output neurons of the retina (*36, 37*). Since the energy required for these processes is thought to be produced by OXPHOS rather than aerobic glycolysis (*3, 38, 39*), energy metabolism in inner retinal neurons and photoreceptors may differ to support their specific needs (*23, 38*).

Because retinal neurons have a high energy demand, any disruption to their sophisticated metabolic system may lead to retinal disease, including retinitis pigmentosa, glaucoma and others (*40, 41*).

Apart from mutations in several metabolic genes, mutations in other genes, such as rhodopsin (*RHO*), are also suspected to contribute to degeneration by indirectly disturbing the cellular energy balance (*40, 42–45*). Therefore, defining the pathways of energy production and consumption is crucial not only for understanding the function of the retina but also for enabling the development of promising therapies for various retinal disorders.

Although numerous studies have investigated energy metabolism in the retina, they lack cellular resolution and do not typically compare inner and outer retinal neurons directly. To fill this knowledge gap, we used two-photon fluorescence lifetime imaging microscopy (2P-FLIM) to quantify the fluorescence lifetime of various metabolic biosensors that we specifically expressed in rods or in inner retinal neurons. FLIM measures the time delay (lifetime) between fluorophore excitation and the emission of a photon (*46*). Because the fluorescence lifetime is an intrinsic property of a given fluorophore, it permits the quantitative assessment of metabolite concentrations (*47*). Moreover, when combined with pharmacological protocols, it provides a powerful approach to investigate the dynamics of cellular energy metabolism. This technique, together with the cell type-specific sensor expression, revealed differential dynamics in the ATP and lactate metabolism between rods and inner retinal neurons in healthy and degenerative retinas.

## Results

### Two photon laser scanning microscopy triggers Ca^2+^ signaling in rod photoreceptors in acute eye sections

The retina is divided into an outer part containing the photoreceptors and an inner part with three types of interneurons and the ganglion cells (*48*). The metabolic ecosystem in the outer retina relies on the exchange of metabolites between RPE cells and photoreceptors, suggesting that the two cell types form a functional unit. In an initial experiment we first visualized the integrity of this unit in our system after the subretinal injection of a mixture of AAVs encoding mCherry under the vitelliform macular dystrophy 2 (VMD2) promoter for RPE-specific expression (*49*) and the calcium sensor GCaMP6s under the control of the mouse opsin promoter (mOP) for rod-specific expression (Fig. 1A) (*26, 50*). Imaging acute eye sections four weeks after the injection confirmed expression of mCherry in RPE cells and GCaMP6s in adjacent rods, revealing the integrity of the RPE/photoreceptor unit (Fig. 1B). Next, we evaluated the functionality of the rods by measuring intracellular Ca^2+^ dynamics in response to light pulses using two-photon laser scanning microscopy (2P) of GCaMP6s (*51*).

**Fig. 1:**
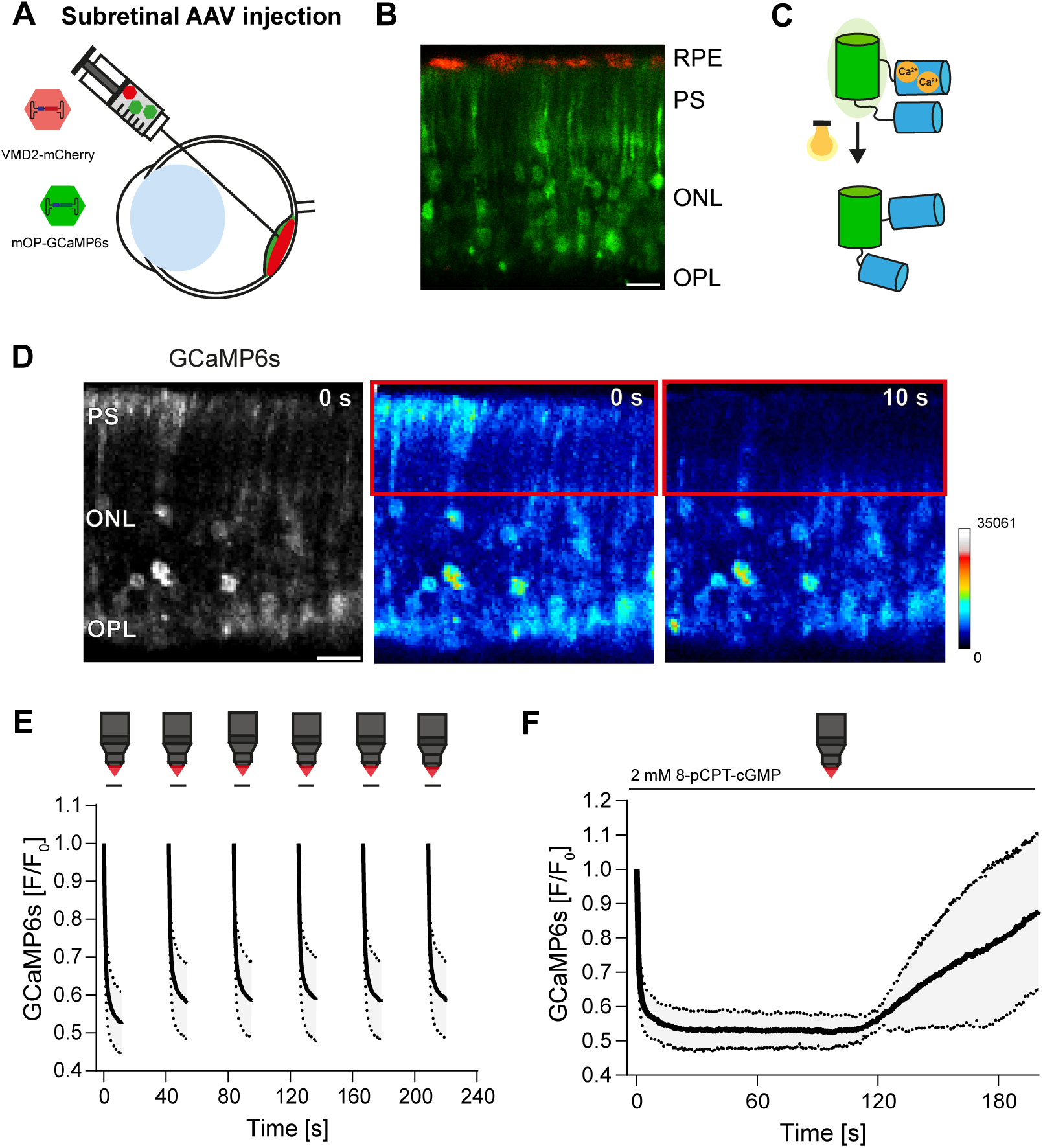
Light-induced calcium response in rods of wildtype mice (BALB/c) during 2P. **A**, Schematic of subretinal injection of a mix containing an AAV encoding mCherry under the VMD2 promoter and an AAV encoding the calcium sensor GCaMP6s under the mOP promoter. **B**, Representative 2P image of an acute eye section from a mouse injected with the AAV mix shown in (A). RPE cells express mCherry (red) and rod photoreceptors GCaMP6s (green). Scale bar: 10 µm. **C**, Schematic of GCaMP6s (adapted from Erofeev *et. al.*(*114*)). Fluorescence intensity of circularly permuted green fluorescent protein (cpGFP) increases when Ca^2+^ is bound. Light-driven reduction in Ca^2+^ in photoreceptors, results in the decrease of fluorescence intensity. **D,** Representative 2P images of rod photoreceptors expressing GCaMP6s. Maximum intensity image at 0 s (left) and intensity weighted images at 0 s (middle) and 10 s (right) of imaging. Red boxes mark the analyzed area (photoreceptor segments). Scale bar: 10 µm. **E,** GCaMP6s traces (mean ± SD) repetitively imaged in photoreceptor segments of wildtype mice (N = 4, n = 3) during 10 s of 2P imaging followed by 30 s recovery. **F,** GCaMP6s traces (mean ± SD) imaged in the photoreceptor segments of wildtype mice (N = 3, n = 3) in the presence of the cGMP analog (2 mM 8-pCPT-cGMP). RPE: retinal pigment epithelium. PS: photoreceptor segments. ONL: outer nuclear layer. OPL: outer plexiform layer.

Activation of the phototransduction cascade by light leads to hydrolysis of cyclic GMP (cGMP), which induces the closure of the cGMP-gated ion channels (CNGCs), decreasing cytosolic Ca^2+^ levels (*52*) and the fluorescence signal intensity of the sensor (Fig. 1C). The GCaMP6s fluorescence in photoreceptor segments was substantially reduced during the 10-second recording interval (Fig. 1D, E), indicating a decrease in intracellular Ca^2+^ levels and the presence of functional, light-responsive rods in our preparation. This response was likely triggered by the light emitted from activated sensors, which falls within the range of the absorption maximum of rhodopsin (*53–56*), or by direct activation through the short infrared light pulses as previously described (*57*). The light-induced drop in calcium was reversed during a 30-second interval in darkness even across multiple imaging and recovery cycles (Fig. 1E), indicating that the system was stable and functional. This was further supported by the increase of Ca^2+^ levels in the presence of the non-cleavable cGMP analog 8-pCPT-cGMP to reopen the channels. As expected, continuous imaging of rod photoreceptor segments over 200 s revealed a slow increase in Ca^2+^ levels during the inflow of the cGMP analog into the chamber (Fig. 1F) (*58, 59*). The delay in the response of approximately 100 seconds corresponds to the time required for 8-pCPT-cGMP to reach the imaging chamber (Fig. S1A). Together, these data verified that rods were fully functional in the acute eye sections and responded to 2P by activating phototransduction (*26, 55*). Importantly, all subsequent data generated by 2P-FLIM, therefore, represent conditions in light-exposed rods.

### Rods rely on glucose whereas inner retinal neurons can use lactate in the absence of glucose to fully maintain ATP levels

To determine the energy source that fuels ATP production in the different retinal neurons, we delivered the ATP sensor ATeam1.03 (*60*) (Fig. 2A) to rods in the outer retina and to ganglion and amacrine cells in the inner retina via subretinal or intravitreal AAV injections (Fig. S1B, C). The fluorescence lifetime (τ_MC_) of the ATP sensor was measured by 2P-FLIM in the presence of glucose or lactate. During the ‘control protocol’ (Table 1), the fluorescence lifetime of ATeam1.03 expressed in rods remained stable for at least 25 min and increased (indicating a reduction in ATP levels) only after removal of glucose and application of NaN_3_ to block OXPHOS (Fig. S2A) (*61*). Importantly, the cells in our sections recovered quickly from the NaN_3_-induced OXPHOS block after a washout period of a few minutes, further confirming the viability of the cells (Fig. S2B).

**Fig. 2:**
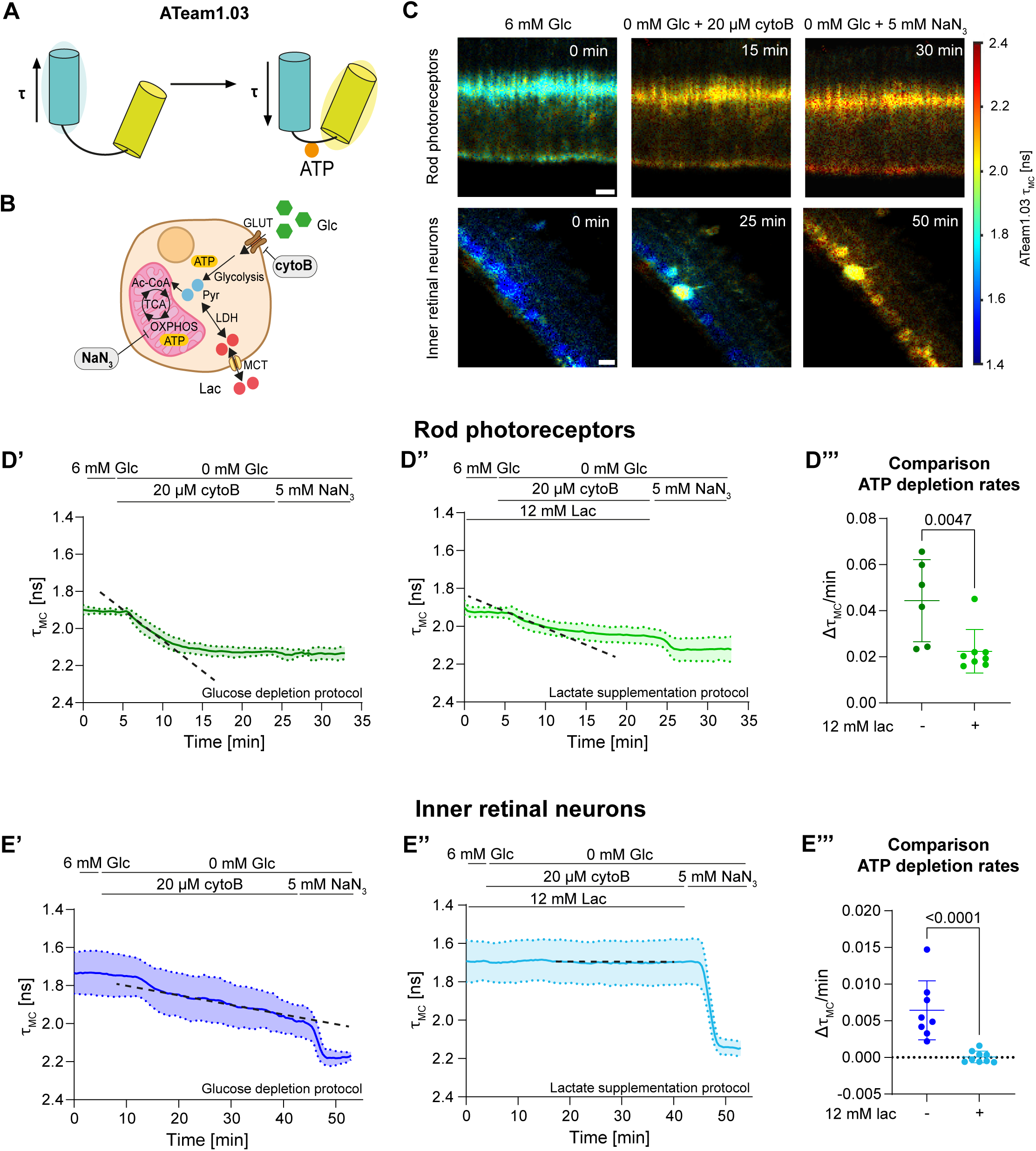
ATP dynamics in rods and inner retinal neurons of wildtype mice (129S6). **A**, Schematic representation of the ATP sensor ATeam1.03 (adapted from Imamura. *et. al.*(*60*)). Binding of ATP decreases the fluorescence lifetime (τ). **B,** Schematic representation of cellular metabolism and the drugs used during the ‘glucose depletion’ and ‘lactate supplementation protocol’: GLUTs were blocked with cytoB and OXPHOS with NaN_3_ (Table 1) during aglycemia. **C,** 2P-FLIM images (τ_MC_) of ATeam1.03 in rods (upper panel) and inner retinal neurons (lower panel) at different time points during the ‘glucose depletion protocol’. Scale bars: 10 µm. **D’, E’,** Lifetime traces of ATeam1.03 in rods (D’; N = 6, n = 6) and inner retinal neurons (E’; N = 8, n = 6) during the ‘glucose depletion protocol’ (mean ± SD). *Black dashed lines*: slopes used to calculate ATP depletion rates. **D’’**, **E’’,** Lifetime traces of ATeam1.03 in rods (D’’; N = 8, n = 5) and inner retinal neurons (E’’; N = 9, n = 5) during the ‘lactate supplementation protocol’ (mean ± SD). *Black dashed lines*: slopes used to calculate ATP depletion rates. **D’’’, E’’’,** Comparison of the ATP depletion rates (Δτ_MC_/min) during the ‘glucose depletion’ (*dark green, dark blue*) and the ‘lactate supplementation protocol’ (*light green, light blue*) in rods (D’’’) and inner retinal neurons (E’’’). Shown are individual values and means ± SD. Mann-Whitney nonparametric test.

**Table 1:** Pharmacological protocols for 2P-FLIM.

| Protocol name | Protocol | Sensor |
| --- | --- | --- |
| Control protocol | 23.5 min ACSF / 6 mM D-glucose → 10 min 0 mM D-glucose + 5 mM NaN<sub>3</sub> | ATeam1.03 |
| NaN<sub>3</sub> outwash protocol | 3.5 min ACSF / 6 mM D-glucose → 10 min 6 mM D-glucose + 5 mM NaN<sub>3</sub> → 16.5 min 6 mM D-glucose | ATeam1.03 |
| Glucose depletion protocol | 3.5 min ACSF / 6 mM D-glucose → 20 (or 40) min 0 mM D-glucose + 20 µM cytoB → 10 min 0 mM D-glucose + 5 mM NaN<sub>3</sub> | ATeam1.03 |
| Lactate supplementation protocol | 3.5 min ACSF / 6 mM D-glucose + 12 mM L-lactate → 20 (or 40) min 0 mM D-glucose + 12 mM L-lactate + 20 µM cytoB → 10 min 0 mM D-glucose + 5 mM NaN<sub>3</sub> | ATeam1.03 |
| Glutamine supplementation protocol | 3.5 min ACSF / 6 mM D-glucose + 6 mM L-glutamine → 20 min 0 mM D-glucose + 6 mM L-glutamine + 20 µM cytoB → 10 min 0 mM D-glucose + 5 mM NaN<sub>3</sub> | ATeam1.03 |
| Lactate transport block protocol | 3.5 min ACSF / 6 mM D-glucose → 40 min 0 mM D-glucose + 20 µM cytoB + 5 µM AR-C155858 → 10 min 0 mM D-glucose + 5 mM NaN<sub>3</sub> | ATeam1.03 |
| Glycogen phosphorylase block protocol | Prior to imaging the sections were pre-incubated for 1 hour with ACSF containing 300 µM DAB / 6 mM D-glucose<br>3.5 min ACSF / 6 mM D-glucose + 300 µM DAB → 40 min 0 mM D-glucose + 20 µM cytoB + 300 µM DAB → 10 min 0 mM D-glucose + 5 mM NaN<sub>3</sub> | ATeam1.03 |
| OXPHOS block followed by aglycemia protocol | 3.5 min ACSF / 6 mM D-glucose → 10 min 6 mM D-glucose + 5 mM NaN<sub>3</sub> → 16.5 min 0 mM D-glucose + 5 mM NaN<sub>3</sub> | ATeam1.03 |
| Lactate production/consumption protocol 1 | 3.5 min ACSF / 6 mM D-glucose → 15 min 6 mM D-glucose + 5 µM AR-C155858 → 15 min 0 mM D-glucose + 5 µM AR-C155858 | LiLac |
| Lactate production/consumption protocol 2 | 3.5 min ACSF / 6 mM D-glucose → 15 min 6 mM D-glucose + 5 µM AR-C155858 + 4 µM Syrosingopine → 15 min 0 mM D-glucose + 5 µM AR-C155858 + 4 µM syrosingopine | LiLac |
| Syrosingopine test protocol | 3.5 min ACSF / 6 mM D-glucose → 15 min 6 mM D-glucose + 4 µM Syrosingopine → 15 min 0 mM D-glucose + 4 µM syrosingopine | LiLac |

We used the ‘glucose depletion protocol’ (Table 1) to investigate changes of intracellular ATP levels in glucose-free conditions. Apart from glucose removal from the media, this protocol included inhibition of glucose transporters (GLUTs) 1, 3 and 4 with cytochalasin B (cytoB) (*62–64*) and subsequent OXPHOS block with NaN_3_ (Fig. 2B). In rods, we observed a fast drop to low steady-state ATP levels in glucose-free conditions (Fig. 2C, D’) and blocking OXPHOS with NaN_3_ did not further reduce ATP levels. In contrast, inner retinal neurons maintained their baseline ATP levels for almost 10 min before ATP levels slowly started to decline in the absence of glucose (Fig. 2C, E’). Importantly, even after 40 min without glucose, inner retinal neurons were not fully depleted of ATP, since blocking OXPHOS further decreased ATP levels (Fig. 2E’). This suggests that inner retinal neurons can produce ATP via OXPHOS over a prolonged period, even if glucose is absent. These data show that rods are highly dependent on glucose and consume ATP faster than inner retinal neurons. The slower drop in ATP levels in the inner retinal neurons suggests that alternative sources support OXPHOS in the absence of glucose.

We investigated if lactate could be such an alternative source to glucose, especially for inner retinal neurons, as it is metabolized via OXPHOS to produce ATP. Therefore, we performed the ‘lactate supplementation protocol’ (Table 1), which included the addition of 12 mM lactate. As before, ATP levels dropped in rods soon after withdrawal of glucose (Fig. 2D’’), albeit with a slower rate (Δτ_MC_/min, Fig. 2D’’’), and ATP remained at intermediate steady-state levels before dropping further after blocking OXPHOS. In sharp contrast, lactate supplementation fully maintained baseline ATP levels in inner retinal neurons for at least 40 min (Fig. 2E’’, 2E’’’) and dropped only after blocking OXPHOS (Fig. 2E’’). This data indicates that inner retinal neurons efficiently produce ATP through OXPHOS when lactate substitutes glucose in the medium. In contrast, rods can only partially maintain ATP levels in the presence of lactate alone. This suggests that rods can use lactate with low efficiency, which is in agreement with the low levels of LDHB in rods (*20, 65*), the enzymatic subunit required for the conversion of lactate to pyruvate that then feeds into OXPHOS. Alternatively, it is also possible that neighboring cells (e.g. Müller glia) converted the lactate in the medium to other metabolites (e.g. glutamine) that were then provided as alternative fuel to photoreceptors, enabling them to partially maintain their ATP levels through OXPHOS (Fig. S3A).

To address this possibility, we replaced glucose by 6 mM glutamine in the medium (the ‘glutamine supplementation protocol’, Table 1). This reduced ATP in rods to intermediate steady-state levels, which dropped further only after blocking OXPHOS (Fig. S3B, C). This shows that rods can indeed use glutamine for ATP production through OXPHOS in the absence of glucose but cannot maintain baseline ATP levels on glutamine alone. Since available inhibitors for glutamine transport or glutaminase were not reliably effective in our system, further investigations with genetic models will be necessary to determine whether some glutamine was provided to rods by other cells during the ‘lactate supplementation protocol’. However, when considered alongside our measurements of lactate levels in rods (see below), it appears unlikely that such a lactate to glutamine conversion shuttle substantially contributed to ATP production under lactate supplemented conditions.

Importantly, these data further underline that rods can use OXPHOS to produce ATP but need glycolysis to fully maintain their ATP levels and thus use a mix of both pathways for energy production.

The rather slow decline of ATP levels in the absence of glucose in inner retinal neurons (Fig. 2E’) suggests the presence of alternative fuels that can be metabolized through OXPHOS in these cells. To test if adjacent cells such as Müller glia cells might release lactate to support glucose-deprived inner retinal neurons, we inhibited monocarboxylate transporter (MCT) 1 and 2 with AR-C155858 (*66*) using our ‘lactate transport block protocol’ (Fig. S4A; Table 1). This did not result in a significant faster ATP decline (Fig. S4B-D) suggesting that monocarboxylates provided by neighboring cells did not significantly impact ATP production in inner retinal neurons. This aligns with the low expression of LDHA in Müller cells (*17*), making the presence of a large lactate pool in these cells unlikely. To test a potential contribution of glycogen, we blocked glycogenolysis by inhibiting the glycogen phosphorylase with 1,4-Dideoxy-1,4-imino-D-arabinitol hydrochloride (DAB, Fig. S4E) in the ‘glycogen phosphorylase block protocol’ (Table 1) (*67*). This resulted in a significantly faster decline of ATP levels (Fig. S4F-H), suggesting that glycogen is an alternative source for ATP in the inner retina. However, we cannot discriminate between the contribution of glycogen internal to inner retinal neurons and glycogen-derived fuel supplied by other cells.

Overall, our data show that rod photoreceptors rely on glucose for ATP production and can only partially maintain ATP levels in the presence of a source such as lactate or glutamine that requires OXPHOS. In contrast, inner retinal neurons can maintain ATP levels equally well in the presence of glucose or lactate. This indicates that rods use both aerobic glycolysis and OXPHOS, while inner retinal neurons use predominantly OXPHOS for ATP production.

### Rods are more glycolytic and inner retinal neurons more oxidative

To directly test if rod photoreceptors rely on both OXPHOS and aerobic glycolysis and inner retinal neurons mostly on OXPHOS, we blocked OXPHOS with NaN_3_ in the presence and absence of glucose (‘OXPHOS block followed by aglycemia protocol’; Table 1). Blocking OXPHOS in the presence of glucose reduced ATP levels in rods to intermediate levels. Only after additionally removing glucose (aglycemia) did ATP levels drop completely (Fig. 3A, B) showing that rods produced ATP via glycolysis but needed OXPHOS to maintain full ATP levels. In contrast, ATP levels in inner retinal neurons declined to very low levels after blocking OXPHOS, and the additional removal of glucose had only a marginal additional effect (Fig. 3A, C). The direct comparison of slope 1 (Fig. 3D) and slope 2 (Fig. 3E) between rods and inner retinal neurons demonstrated significant differences in ATP production: while rods produced ATP via both aerobic glycolysis and OXPHOS, inner retinal neurons rely almost exclusively on OXPHOS.

**Fig. 3:**
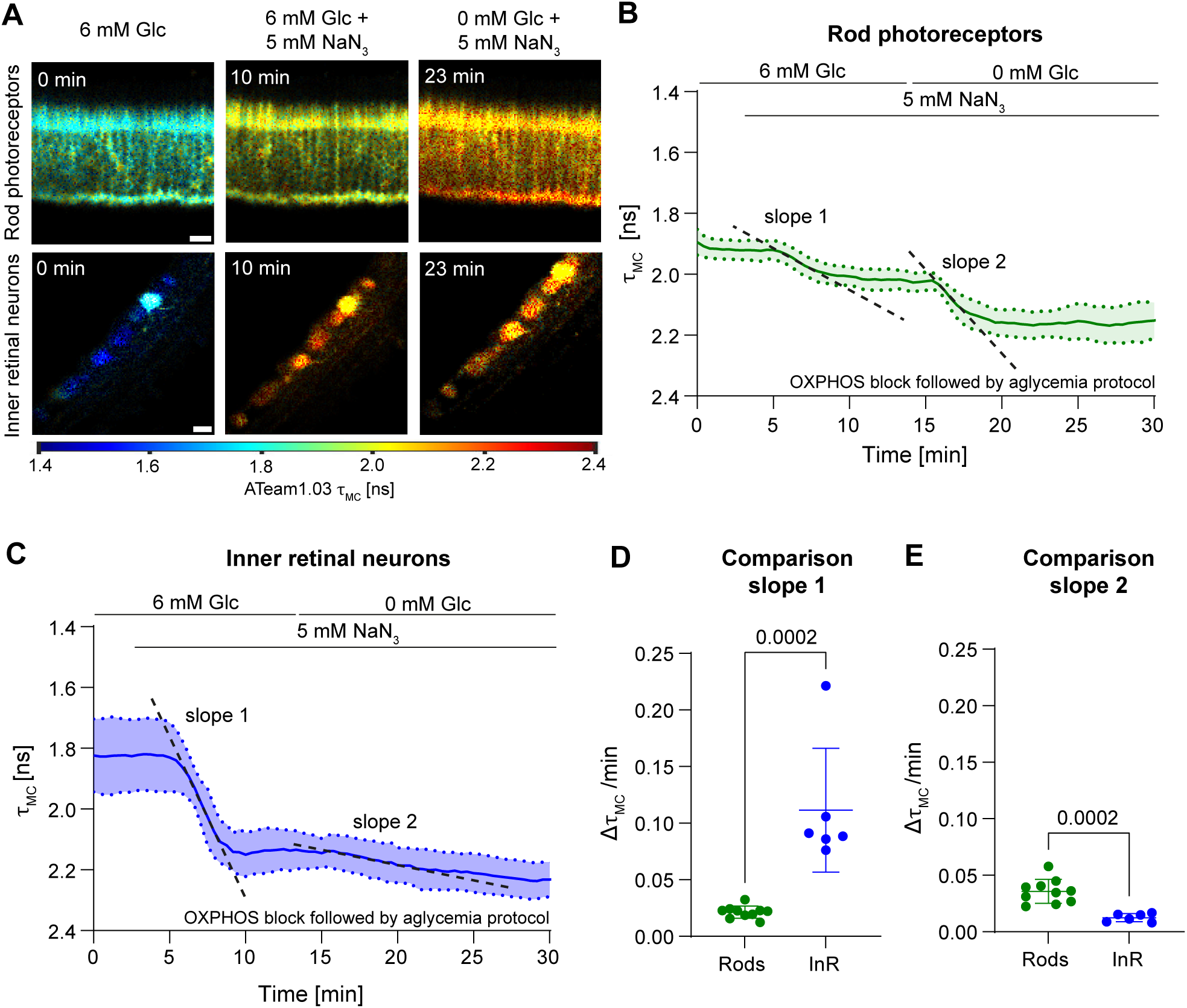
ATP production via aerobic glycolysis versus OXPHOS in rods and inner retinal neurons of wildtype mice (129S6). **A,** 2P-FLIM images (τ_MC_) of ATeam1.03 in rods (upper panel) and inner retinal neurons (lower panel) at different time points during the ‘OXPHOS block followed by aglycemia protocol’ (Table 1). Scale bars: 10 µm. **B, C**, Lifetime traces of ATeam1.03 in rods (B; N = 11, n = 8) and inner retinal neurons (C; N = 6; n = 5) during the OXPHOS block with NaN_3_ followed by aglycemia (0 mM Glc; mean ± SD). *Black dashed lines*: slopes used to calculate ATP depletion rates. **D, E,** Comparison of ATP depletion rates (Δτ_MC_/min) in rods (*green*) and inner retinal neurons (*blue*) after OXPHOS block with NaN_3_ (slope 1, data from Fig. 3B, C) and subsequent glucose removal (slope 2, data from Fig. 3B, C). Shown are individual values and means ± SD. Mann-Whitney nonparametric test.

To further validate that inner retinal neurons are more oxidative than rods, we expressed Peredox, a fluorescent biosensor that reports the cytosolic NADH/NAD^+^ ratio and thus the redox state of the cell (Fig. S5A) (*68, 69*). Cytosolic NADH is mainly produced by glyceraldehyde 3-phosphate dehydrogenase (GAPDH) which links the cytosolic NAD(H) pool with glycolysis. For glycolysis to proceed, NADH has to be re-oxidized to NAD^+^, which is mainly done by mitochondrial shuttles.

However, in glycolytic cells the high glycolytic rate results in saturation of mitochondrial shuttles, and regeneration of NAD^+^ via LDHA is necessary (Fig. S5B) (*70*). Consequently, glycolytic cells have a higher cytosolic NADH/NAD^+^ ratio than oxidative cells. A low fluorescence lifetime of Peredox therefore indicates an oxidative state, and a high fluorescence lifetime a glycolytic state of the cell (*69*). The higher fluorescence lifetime of Peredox in rods at baseline (Fig. S5C, D) supports our conclusions from the ATP sensor that photoreceptors favor aerobic glycolysis, whereas inner retinal neurons rely more on mitochondrial respiration for energy production.

### Rods produce more lactate than inner retinal neurons

A century ago, Otto Warburg showed that the retina produces lactate and concluded that some retinal cells must perform aerobic glycolysis (Warburg effect) to generate ATP (*10*). Since our data indicate that rods are the main cells using aerobic glycolysis, we sought to determine if the retinal neurons differ in their production of lactate, the final product of aerobic glycolysis. To this end, we used LiLac, a single fluorophore biosensor that decreases its fluorescent lifetime upon binding lactate (Fig. 4A) (*71*). To measure lactate production and consumption in the retinal cells, we added AR-C155858 (*66*) that blocks MCT1 and MCT2, the main lactate transporters in photoreceptors and inner retinal neurons (*72*). Blocking MCT1/2 in the presence of glucose (the ‘lactate production/consumption protocol’, Fig. 4B; Table 1) resulted in higher intracellular lactate levels in rods than in inner retinal neurons. Moreover, the rate of lactate accumulation was significantly higher in rods (Fig. 4C-E; slope 1), further supporting our conclusion that rods have a high capacity to perform aerobic glycolysis. After removal of glucose, lactate levels rapidly declined (slope 2; Fig. 4D, F) in both rod photoreceptors and inner retina neurons, suggesting that both rods and inner retinal neurons can metabolize lactate. Since rods express only low levels of LDHB, the enzymatic subunit that is required for an efficient conversion of lactate into pyruvate for OXPHOS, the strong decline of lactate in rods in the presence of the MCT1/MCT2 blocker AR-C155858 was surprising and prompted us to test if lactate could have ‘leaked’ to the extracellular space through MCT4 that has been reported to be present in rods, albeit only at low levels (*72*). However, the addition of the MCT1/4 blocker syrosingopine (*27, 73*) together with the MCT1/2 blocker during the ‘lactate production/consumption protocol 2’ (Fig. S6A, Table 1) did not change lactate accumulation and reduction in rods compared to when only MCT1 and MCT2 were blocked (Fig. S6B-E). The minimal contribution of MCT4 to lactate exchange between rods and the extracellular environment was further evidenced when syrosingopine, which has a 60-fold higher potency for MCT4 than for MCT1 (*73*), was used alone during the ‘syrosingopine test protocol’ (Table 1). Here, we observed a much slower and delayed increase of intracellular lactate compared to the block of MCT1/2 transporters (Fig. S6F, G). This showed that the MCT1/4 blocker was functional and supported the conclusion that MCT4 is not a major transporter for lactate in rods. When glucose was withdrawn in the presence of syrosingopine, however, lactate declined with a rate similar to that observed during the MCT1/2 block protocol (Fig. S6H), again suggesting that rods metabolized lactate. In summary, this data indicates that rods produce lactate in the presence of glucose but can also consume lactate in the absence of glucose, despite the low expression level of LDHB. This data is consistent with our findings in Fig. 2D’’. Inner retinal neurons are rather lactate consumers, consistent with the data in Fig. 2E’’, producing less lactate and at a lower rate compared to rods, but also consume lactate in the absence of glucose.

**Fig. 4:**
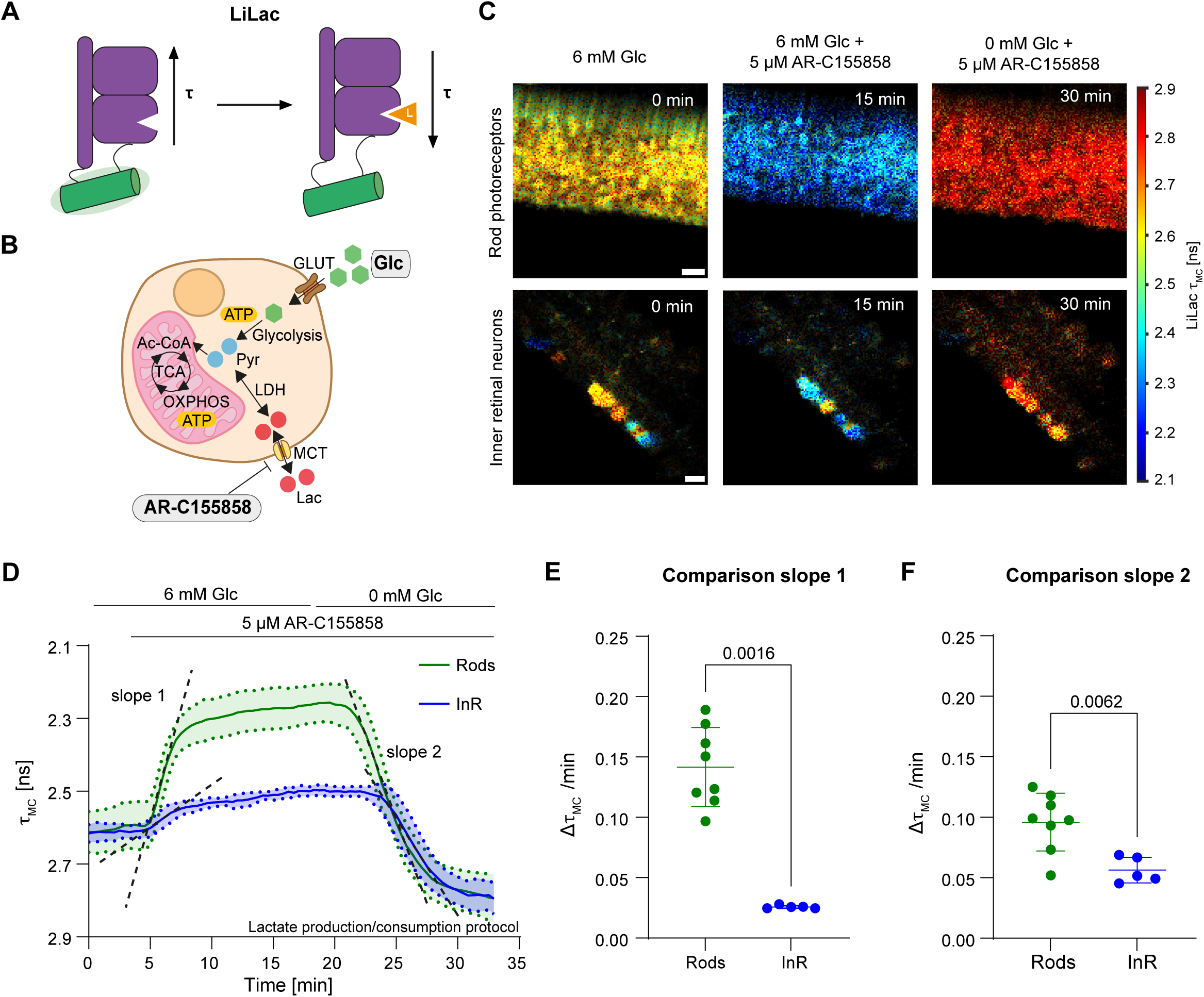
Lactate production and consumption in rods and inner retinal neurons of wildtype mice (129S6). **A,** Schematic representation of the lactate sensor LiLac (adapted from Koveal, *et. al.*(*71*)). The lifetime (τ) of LiLac decreases when bound to lactate (L). **B,** Schematic representation of cellular metabolism, illustrating lactate transport via MCTs and their blocking with AR-C155858. **C,** 2P-FLIM images (τ_MC_) of LiLac in rods (upper panel) and inner retinal neurons (lower panel) at different time points during the ‘lactate production/consumption protocol’ (Table 1). Scale bars: 10 µm. **D,** Lifetime traces (means ± SD) of LiLac in rods (*green*; N = 8; n = 3) and inner retinal neurons (*blue*; N = 5, n = 3) during the ‘lactate production/consumption protocol’. *Black dashed lines*: slopes used to calculate the changes in the lifetime in LiLac shown in panels E, F. **E, F**, Lifetime changes (Δτ_MC_/min) in LiLac during the MCT block in the presence of glucose (E; slope 1; lactate production) and in the absence of glucose (F; slope 2; lactate consumption) in rods (*green*) and inner retinal neurons (*blue*). Shown are individual values and means ± SD. Mann-Whitney nonparametric test.

### Energy metabolism in degenerating rods of *Rho^P23H/+^* mice

To investigate metabolic changes in response to inherited retinal degeneration, we repeated the experiments in rods of *Rho^P23H/+^* mice, an established model for retinitis pigmentosa (RP) (*74*). In this model, mutant rhodopsin accumulates in the endoplasmic reticulum (ER), thereby triggering the unfolded protein response (UPR). This causes ER stress and subsequently leads to retinal degeneration (*44, 75, 76*). The loss of rod photoreceptors may reduce the supply of rod-derived lactate to RPE cells, which might then be forced to utilize glucose from the choriocapillaris for their own needs, instead of passing it on to the photoreceptors, thereby potentially disrupting the metabolic ecosystem (*45, 77*).

To measure metabolite dynamics in rods of the degenerating retina, we used *Rho^P23H/+^* mice at 8 weeks of age when the thickness of the outer nuclear layer was already reduced by approximately 50% and photoreceptor segments were disorganized and shortened (Fig. S7A) (*74*). To confidently compare the *Rho^P23H/+^* data to wildtype rods, it was important to first evaluate the vital state of the degenerating rods at this age by measuring Ca^2+^ in response to imaging using GCaMP6s (Fig. S7B). As observed in wildtype cells (Fig. 1), the Ca^2+^ levels dropped in photoreceptor segments of the *Rho^P23H/+^*mouse during imaging and recovered within 30 s in darkness (Fig. S7C, D). GCaMP6s measurement after inflow of 8-pCPT-cGMP showed that Ca^2+^ levels started increasing after the cGMP analog reached the imaging chamber, suggesting that the cGMP-gated ion channels reopened (Fig. S7E).

Therefore, rods in 8-week-old *Rho^P23H/+^* mice were in a light-activated state during imaging and exhibited functional properties similar to those of wildtype rods. This enabled us to directly compare the metabolic dynamics of healthy and diseased photoreceptors.

To determine whether degenerating processes changed the ATP dynamics in rods, we applied the ‘glucose depletion protocol‘ and ‘the lactate supplementation protocol‘ (Table 1). We observed that the drop of ATP levels in the absence of glucose was similarly rapid in degenerating and healthy rods (Fig. 5A-C). Like healthy rods, degenerating rods could partially maintain ATP levels when lactate substituted glucose (Fig. 5D). The decline in ATP levels in the presence of lactate was slower compared to that in the ‘glucose depletion protocol’ (Fig. 5E) and it was faster than in wildtype rods (Fig. 5F). This suggests that ATP production via OXPHOS is less efficient in degenerating than wildtype rods in the presence of lactate.

**Fig. 5:**
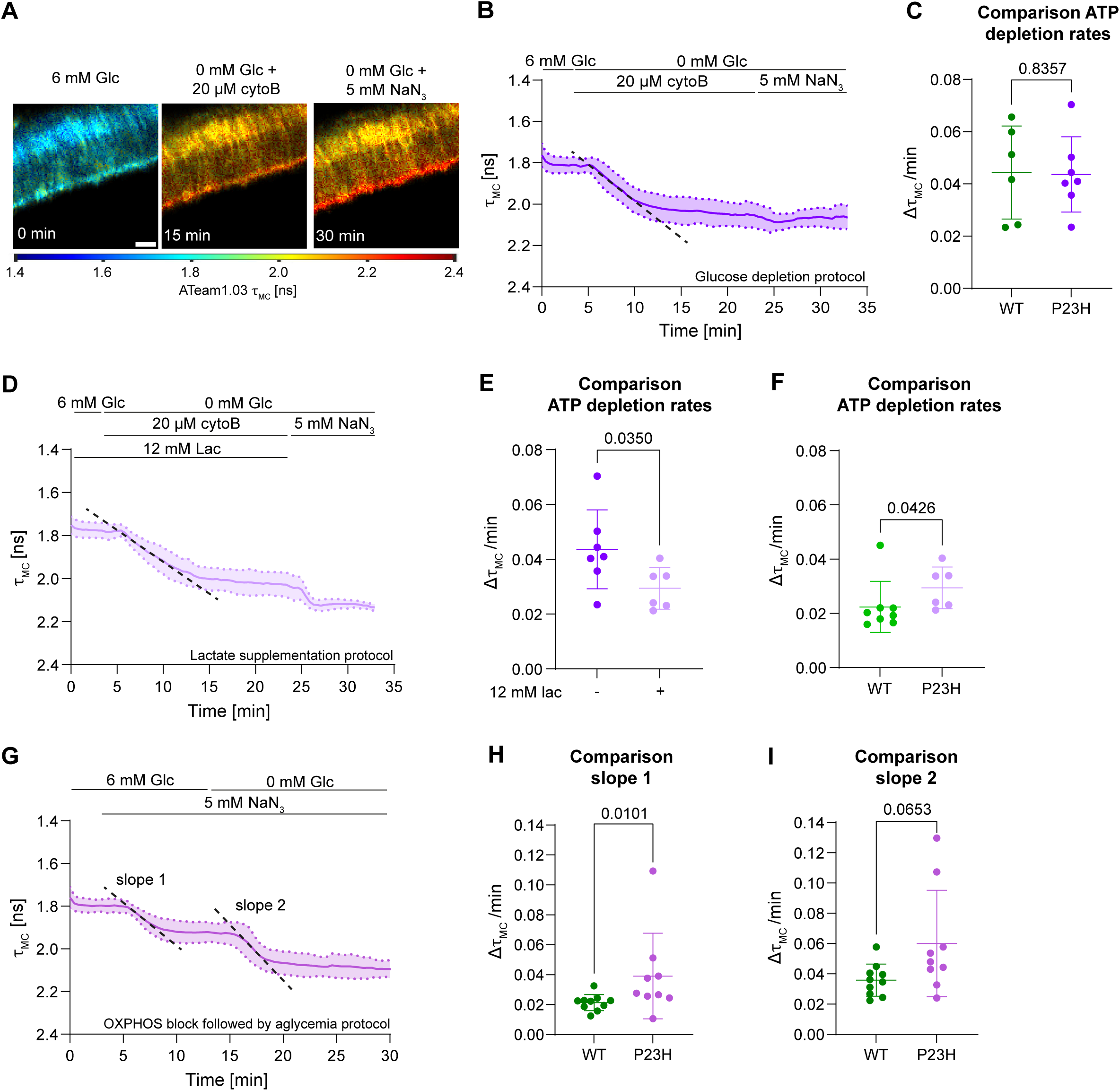
ATP dynamics in degenerating rods of *Rho^P23H/+^* mice. **A,** 2P-FLIM images (τ_MC_) of ATeam1.03 in rods of *Rho^P23H/+^* mice at different time points during the ‘glucose depletion protocol’ (Table 1) and **B,** corresponding lifetime trace (mean ± SD; N = 7, n = 5). Scale bar: 10 µm. *Black dashed line*: slope used to calculate ATP depletion rates. **C,** Comparison ATP depletion rates (Δτ_MC_/min) during the ‘glucose depletion protocol’ in rods of wt (*green*; data from Fig. 2D’’’) and *Rho^P23H/+^* mice (*violet*). Shown are individual values and means ± SD. Mann-Whitney nonparametric test. **D,** Lifetime trace of ATeam1.03 in rods of *Rho^P23H/+^* mice during the ‘lactate supplementation protocol’ (Table 1; N = 6, n = 5). *Black dashed line*: slope used to calculate the ATP depletion rate. **E,** Comparison of ATP depletion rates (Δτ_MC_/min) during the ‘glucose depletion’ (*dark violet*) and the ‘lactate supplementation protocol’ (*light violet*). Shown are individual values and means ± SD. Mann-Whitney nonparametric test. **F,** Comparison of ATP depletion rates (Δτ_MC_/min) in wt (*green*; data from Fig. 2D’’’) and *Rho^P23H/+^* mice (*light violet*) during the ‘lactate supplementation protocol’. Shown are individual values and means ± SD. Mann-Whitney nonparametric test. **G,** Lifetime trace of ATeam1.03 in rods of *Rho^P23H/+^* mice during the ‘OXPHOS block followed by aglycemia protocol’ (Table 1; N = 9, n = 5). *Black dashed lines*: slopes used to calculate ATP depletion rates. **H, I,** Comparison of ATP depletion rates (Δτ_MC_/min) in wt (*green*; data from Fig. 3D, E) and *Rho^P23H/+^* mice (*violet*) during the OXPHOS block (H; slope 1) followed by aglycemia (I; slope 2) protocol. Shown are individual values and means ± SD. Mann-Whitney nonparametric test.

Degenerating rods, as wildtype rods, performed both aerobic glycolysis and OXPHOS as indicated by the two-step reduction in ATP levels after OXPHOS block and additional glucose withdrawal (Fig. 5G). However, the initial reduction (slope 1; Fig. 5G, H) was steeper in degenerating rods, which suggests that blocking OXPHOS had a greater impact on ATP levels in degenerating rods than in healthy ones. However, the contribution of aerobic glycolysis to ATP levels was similar in wildtype and degenerating rods (Fig. 5I).

### Degenerating rods can produce and likely also consume lactate

Our results indicated subtle metabolic changes in the use of lactate for ATP production in degenerative rods. To directly test lactate production and consumption, we expressed LiLac in rods of the *Rho^P23H/+^* mouse. Since our results from wildtype mice indicated that MCT1 and 2 but not MCT4 are the predominant transporters responsible for lactate transport in photoreceptors, we used the MCT1/2 blocker to measure lactate production and consumption in degenerating rods. This data showed that degenerating rods accumulated more lactate in the presence of the blocker, and did so at a faster rate than wildtype rods (Fig. 6A-C; slope 1). The rate of lactate reduction in the absence of glucose was similar in normal and degenerating rods (Fig. 6B, D; slope 2).

**Fig. 6:**
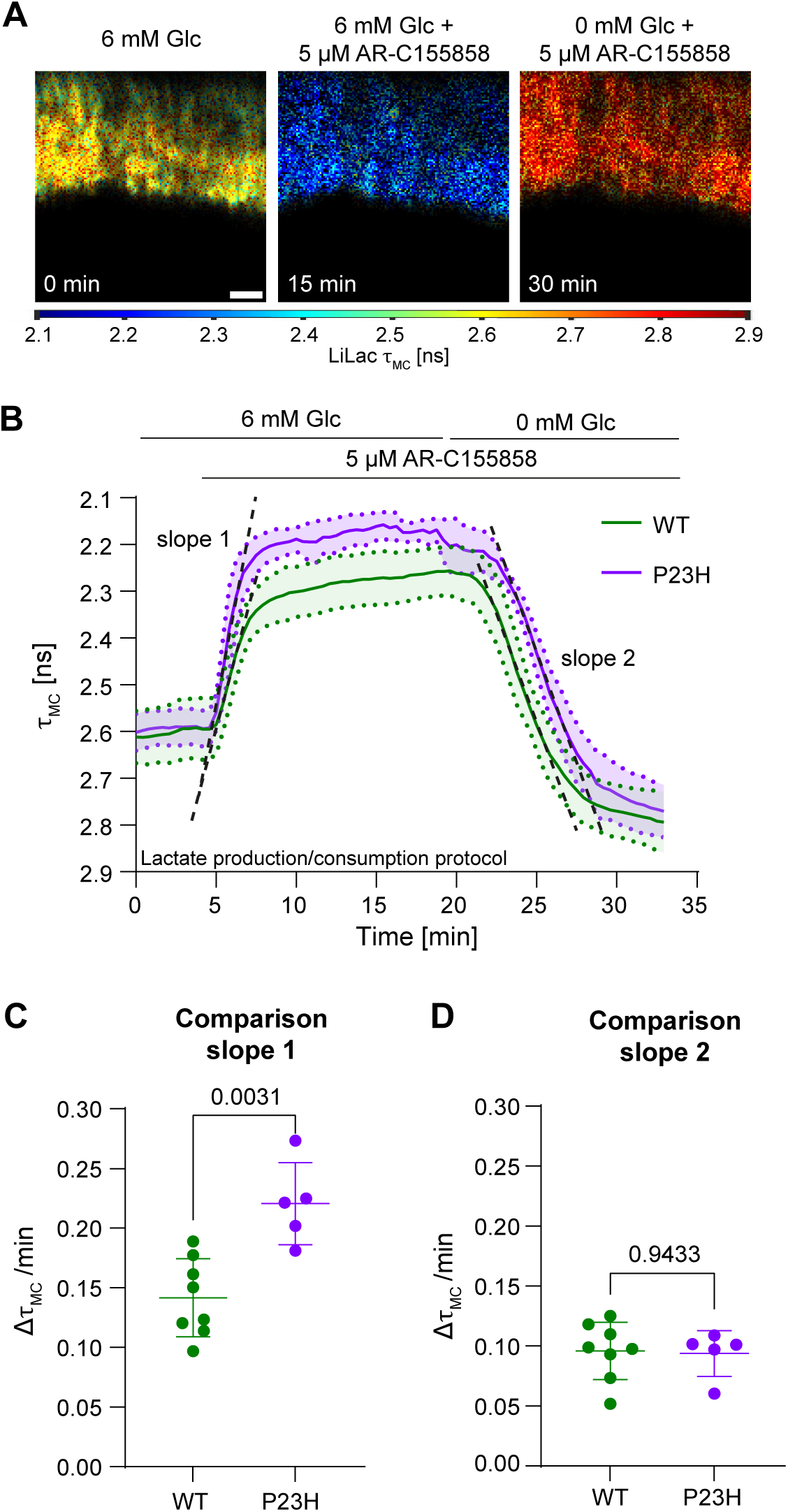
Lactate production in rods of *Rho^P23H/+^* mice. **A,** 2P-FLIM images (τ_MC_) of the lactate sensor LiLac in rods of *Rho^P23H/+^* mice during the ‘lactate production / consumption protocol’ (Table 1). Scale bar: 10 µm. **B,** Lifetime traces (means ± SD) of LiLac in rods of wt (*green*; data from Fig. 4D; N = 8, n = 3) and *Rho^P23H/+^*mice (*violet*; N = 5, n = 4) during the ‘lactate production / consumption protocol’ (Table 1). *Black dashed lines*: slopes used to calculate the lifetime changes of LiLac in wt and *Rho^P23H/+^*mice. **C, D,** Lifetime changes (Δτ_MC_/min) in LiLac during the MCT block in the presence of glucose (C; slope 1; lactate production) and in the absence of glucose (D; slope 2; lactate consumption) in rods of wt (*green*; data from Fig. 4E, F) and *Rho^P23H/+^* mice (*violet*). Shown are individual values and means ± SD. Mann-Whitney nonparametric test.

Overall, our data show that rods of the *Rho^P23H/+^* mouse, like rods of wildtype mice, rely on glucose as energy source and that an alternative fuel that supports ATP production through OXPHOS was not sufficient to maintain ATP levels. Since lactate production was slightly increased, *Rho^P23H/+^* rods might have shifted towards aerobic glycolysis.

## Discussion

A century ago, Otto Warburg reported that the retina produces lactate, which indicated that some retinal cells use aerobic glycolysis to produce energy (*10*). Although many subsequent studies have confirmed this observation and have shown that photoreceptors express key enzymes involved in aerobic glycolysis, such as LDHA, PKM2, and HK2, they did not directly compare metabolic dynamics between different retinal cell types (*13–21*). To address this, we expressed genetically encoded metabolic biosensors in rod photoreceptors and inner retinal neurons and recorded the cell type-specific responses to pharmacological protocols with 2P-FLIM. Using this approach, we collected several data sets that demonstrated, among others, that rods are glycolytic (but also use OXPHOS) and inner retinal neurons are oxidative: rods had a higher NADH/NAD^+^ ratio (Fig. S5), were unable to maintain ATP at baseline levels in the presence of lactate or glutamine, metabolites that contribute to ATP production via OXPHOS (Fig. 2, S3), produced more lactate and did so at a higher rate (Fig. 4), and showed a lower ATP depletion rate in the absence of OXPHOS (Fig. 3).

OXPHOS generates thirty-six molecules of ATP from one molecule of glucose, whereas glycolysis generates only two ATPs (*78*). Therefore, using aerobic glycolysis may appear to be an inefficient strategy for rods that require vast amounts of energy for function. However, glycolysis is more rapid than OXPHOS and produces a large number of carbon intermediates that can be used to produce the building blocks needed to satisfy the anabolic needs of the cells (*30, 78*). Given that photoreceptors shed and renew about 10% of their outer segments daily (*79*), they have an exceedingly high biomass turnover rate, requiring large amounts of lipids, amino acids, and nucleotides to maintain their cellular integrity and function, which justifies their reliance on aerobic glycolysis (*13*). However, our data clearly show that rods metabolize glucose not only by aerobic glycolysis but simultaneously also by OXPHOS. Although rods use both pathways to maintain optimal ATP levels and sustain their anabolic and metabolic demand, they can survive for several months without *Cox10* (*26*), an essential assembly factor for complex IV of the mitochondrial electron transport chain central to OXPHOS, or without LDHA (*13, 17*), the key enzyme that converts pyruvate to the glycolytic end product lactate.

We speculate that rods can cope for some time with the imbalance between aerobic glycolysis and OXPHOS when relying on either pathway alone but eventually degenerate because they cannot tolerate the changes in ATP production and/or anabolic processes long-term.

If rods can cope without OXPHOS for months, why do they have such a massive load of mitochondria? *Ex vivo* studies showed that photoreceptors produce succinate that is converted into malate by the RPE, which in turn can refuel succinate production in the retina via reverse succinate dehydrogenase (SDH) activity in the TCA cycle (*80*). This may be beneficial for the anabolism in photoreceptors because it serves as an anaplerotic pathway to maintain metabolic pools (*81*). Since the reverse SDH reaction diverts electrons out of the electron transport chain (*80*), it uncouples mitochondria, leading to reduced ATP production via OXPHOS. This may help to explain the presence of the large number of mitochondria in photoreceptors and suggests that rods may depend more on a functional TCA cycle than on efficient ATP production via OXPHOS (*26*).

The failure of lactate and glutamine to fully substitute for glucose (Fig. 2, S3) and the partial reduction in ATP levels in the absence of OXPHOS (Fig. 3) indicate that glucose is the main energy substrate in rods, as suggested before (*32, 82*). This aligns with the notion that photoreceptors express low levels of LDHB but high levels of LDHA (*13, 20*), favoring lactate production over lactate consumption. Nevertheless, the partial maintenance of ATP levels in the presence of lactate instead of glucose (Fig. 2) suggests that rods can convert lactate to pyruvate and use it in OXPHOS despite the low intrinsic LDHB level (*20, 65*). The potential of rods to consume lactate is further supported by the rapid decline of intracellular lactate levels in the absence of glucose and presence of MCT blockers (Fig. 4, S6). However, the partial maintenance of ATP levels in the presence of glutamine (Fig. S3) may provide an alternative explanation for ATP production when lactate is the only fuel available. It is possible that neighboring cells could have metabolized lactate from the medium into alternative fuels such as glutamine and provided them to rods (*19, 40*). Nevertheless, the fast decline of lactate levels in presence of all MCT blockers (Fig. S6C-E) indicates that rods indeed can metabolize lactate, at least to some extent. Although rods can use alternative fuels for ATP production through OXPHOS, neither lactate nor glutamine supplementation resulted in ATP levels that were reached in the presence of glucose. Together with the reduced steady-state ATP levels in the absence of OXPHOS, this clearly indicates that rods need both glycolysis and OXPHOS to sustain full energy levels.

In sharp contrast to rods, inner retinal neurons fully maintained their ATP levels when lactate instead of glucose was provided in the medium, indicating that inner retinal neurons metabolize lactate and rely on OXPHOS for ATP production (Fig. 2). This data also aligns with data showing that inner retinal neurons express high levels of LDHB (*17*), which favors lactate consumption over lactate production.

While ATP levels in rods dropped quickly in the absence of glucose, those in inner retinal neurons diminished more slowly (Fig. 2). This suggests that either their ATP consumption was lower or that they used alternative fuels, such as lactate, for energy production through OXPHOS, as suggested by others (*27*). In analogy to the astrocyte-neuron lactate shuttle in the brain (*83, 84*), lactate could be supplied by neighboring cells, including Müller glia, although clear evidence for this is missing (*85*). However, since blocking lactate transport did not affect the ATP depletion rate in the absence of glucose, our data does not favor the presence of a lactate shuttle from Müller cells to inner retinal neurons. This might be expected, as Müller cells express low levels of LDHA (*17*) and are deficient for all PKM isoforms (*19*) making an efficient production of lactate in these cells unlikely. Furthermore, blocking glycogenolysis in the absence of glucose significantly increased the ATP depletion rate in inner retinal cells (Fig. S4), even though lactate transport was not blocked. This additionally challenges the presence of a lactate shuttle to inner retinal neurons and instead suggests that inner retinal neurons can use glucose-6-phosphate, the breakdown product of glycogen, to produce some of their ATP through OXPHOS. Whether glycogenolysis occurs in the neurons or in adjacent cells still needs to be defined. As neither blocking lactate transport nor inhibiting glycogen breakdown accelerated ATP depletion in inner retinal neurons to a rate similar to that observed in glucose-starved rods, our data support the notion that photoreceptors have the highest energy consumption of all retinal cells.

We need to emphasize that by expressing the biosensors under control of the human synapsin 1 (hSyn1) promoter fragment (*86*), we analyzed metabolites in a heterogeneous cell population of ganglion and amacrine cells (Fig. S1C) that may differ in their metabolic properties (*23*). Furthermore, we cannot discriminate between the different subtypes of these neurons nor between individual cellular compartments, which may differ in their mitochondrial distribution and local energy metabolism (*3, 87*). Nevertheless, our results clearly demonstrate that ganglion and amacrine cells in the inner retina mostly rely on OXPHOS for ATP production, whereas rods in the outer retina perform both aerobic glycolysis and OXPHOS.

Our data obtained from wildtype mice support the proposed role of rod photoreceptors in the metabolic landscape and demonstrate that glucose is the primary substrate for energy and lactate production through aerobic glycolysis in rods. To investigate if the metabolic properties of photoreceptors are altered during degeneration, we used the *Rho^P23H/+^*mouse, a well-established model of RP (*74*). Perhaps surprisingly, our data revealed only minor metabolic alterations in the degenerating rods. However, despite the comparable ATP depletion rates in the absence of glucose and alternative fuels (Fig. 5C), ATP levels decreased faster in *Rho^P23H/+^* rods when only lactate was supplied to the cells (Fig. 5F). This suggests that lactate usage and/or ATP production via OXPHOS may be impaired in the disease model. Indeed, the accumulation of mutant rhodopsin in the ER is linked to oxidative stress, which can cause mitochondrial dysfunction (*44, 75, 76, 88–91*). Reduced mitochondrial function in *Rho^P23H/+^* rods may require an increase in glycolysis and LDH activity (*92, 93*) as a compensatory response resulting in heightened lactate production, as our data indicated (Fig. 6). However, this may not fully compensate for the lactate that is no longer produced due to the loss of rods during degeneration. In consequence, less lactate would be available for RPE cells, causing them to use more glucose, which in turn may impact metabolism in the remaining photoreceptors (*77, 94*). However, we only observed a minor metabolic effect in our *Rho^P23H/+^*rods. This may be due to the availability of abundant metabolites in the medium during 2P-FLIM, which contrasts the reduced glucose abundance due to the restricted transport in the *Rho^P23H/+^* retina *in vivo*. Importantly, however, our data indicate that the mutant rods maintain their capability to efficiently metabolize glucose, like cones that can be activated by the subretinal injection of glucose in the degenerative *Rho^P23H/+^*retina (*94*). Targeting energy metabolism might therefore be a promising therapeutic approach to restore mitochondrial function in both inherited and acquired diseases, as also suggested by others (*40, 45, 95–97*).

Previous studies used enzymatic stainings, genetic models, metabolic flux analysis of whole retinas, proteomics, or single-cell RNA transcriptomics to study retinal metabolism (*13, 17, 19, 23, 43, 98–101*). However, 2P-FLIM provides a higher temporal and spatial resolution that has allowed us to observe differences in metabolic dynamics in the different cell types of the retina. 2P-FLIM also overcomes the limitations of ratiometric biosensors for the quantitative assessment of metabolite concentrations (*47*) as it is insensitive to differences in sensor expression levels.

A limitation of our study is that the *ex vivo* approach does not mimic the regulated delivery of metabolites and oxygen to the retina through the blood/retina barrier (*12, 102*). The use of retinal slices results in a uniform and unrestricted access of the cells to the metabolites in the medium, which differs from the *in vivo* situation where the distance from the supplying blood vessels defines a concentration gradient for the nutrients in the tissue, affecting the supply for the cells. Further, the cells in the *in vivo* retina are simultaneously exposed to different metabolites in variable concentrations, which may have an influence on energy production. Still, by providing distinct metabolites in the medium, we clearly defined the capabilities of the individual cell types to metabolize specific fuels for ATP production and showed how cells depend on different energy pathways to maintain their ATP levels. Additional genetic experiments, including cell-type-specific knockouts of individual transporters, will nevertheless be needed to completely dissect the metabolic interaction between neighboring cells *in vivo*.

Overall, the versatility of genetically encoded metabolic biosensors combined with 2P-FLIM and pharmacological protocols enables the dissection of the metabolism on a cellular level, which further improves our understanding of the metabolic landscape in the retina. Our data demonstrate, at a cellular resolution, that rod photoreceptors, but not inner retina neurons, perform aerobic glycolysis but also need OXPHOS to maintain ATP levels. We show that rods can use glutamine as an alternative fuel and that rods can metabolize lactate even though they express only low levels of LDHB. Further, our data show that a mutation in rhodopsin induces only subtle changes in rod-specific energy metabolism. Our study complements and extends results generated by other methods by increasing the spatial and temporal resolution. The advantages offered by 2P-FLIM will enable us to test if rods and cones differ in their metabolism, investigate metabolic dynamics in the RPE, and address night-day differences in the energy production in photoreceptors in the near future.

## Methods

### Study design

The objective of this study was to dissect the metabolic differences between rod photoreceptors and inner retinal neurons, as well as to determine whether rod photoreceptor metabolism is impaired in the *Rho^P23H/+^* mouse, a model of RP. To achieve this, we express genetically encoded biosensors under the control of cell type-specific promoters using AAV-mediated gene transfer to the retinal neurons. To determine the intracellular levels of ATP and lactate, as well as the redox status of the cells, and to resolve the metabolic dynamics of the different neurons, we applied a variety of pharmacological protocols and measured the fluorescence lifetime of the biosensors using 2P-FLIM. All animal experiments and animal care were conducted in accordance with institutional regulations of the University of Zürich and the regulations of the veterinary authorities of the Kanton of Zürich.

### Animals

All animal experiments were approved by the Cantonal Veterinary Office of Zurich, Switzerland (license numbers: ZH019/2019, ZH091/2019, ZH105/2022, ZH065/2025) and adhered to the ARVO statement for the use of Animals in Ophthalmic and Vision Research. Wildtype (wt) mice (129S6, C57B6/J) and transgenic mice (*Rho^P23H/+^*) (*74*) on a C57B6/J background were maintained as breeding colonies at the Laboratory Animal Services Center (LASC) of the University of Zurich with a 14/10 h light-dark cycle with an average light intensity of 60-150 lux at cage level. BALB/c mice were obtained from Charles River (Hilden, Germany). Mice had access to food and water *ad libitum*.

### Cloning and virus production

The cDNAs encoding the biosensors for calcium (GCaMP6s (*103*)), ATP (ATeam1.03 (*60*), (Viral Vector Facility of the Neuroscience Center Zurich; VVF)), lactate (LiLac (*71*) (Addgene #184570)) and NADH/NAD^+^ (Peredox (*68*); VVF) were cloned behind the rod-specific mOP promoter (*26, 50*) and the hSyn1 promoter (*86*). To avoid recombination during AAV production, a codon-diversified version of ATeam1.03 was used (*104*). The construct containing GCaMP6s was packaged into the AAV8(double Y-F + TV)/2 capsid (*26, 105*). Constructs containing ATeam1.03, LiLac and Peredox were packaged into the AAV2(QuadYF+TV; 7m8)/2 capsid (*106*) for transduction of rods after subretinal injection, or the AAV6 capsid for transduction of inner retinal neurons after intravitreal injection. To label RPE cells the mCherry cDNA was cloned behind the VMD2 promoter (*49*) and packaged into the AAV4 capsid. All AAVs were produced by the VVF of the Neuroscience Center Zurich.

### Intraocular injection of AAVs and funduscopy

Intraocular injections were performed as described previously (*107*). Briefly, prior to injection pupils of the mice were dilated under dim light with 1 drop of 1% Cyclogyl (Alcon Pharmaceuticals, Fribourg, Switzerland) and 1 drop of 5% Neosynephrin (Ursapharm Schweiz GmbH, Hünenberg, Switzerland). The mice were anesthetized under dim light with a single subcutaneous injection of Ketamine (85 mg/kg, Pfizer AG, Zürich, Switzerland) / Xylazine (10 mg/kg, Elanco Animal Health GmbH, Basel, Switzerland) using a syringe with a 30G needle. Lacrinorm Carbomerum (Bausch & Laumb Swiss AG, Zug, Switzerland) was applied to keep the eyes moist. Anesthetized mice were placed on a heating pad set at 37°C and the head was fixed in a stereotactic adapter (Hugo Sachs Elektronik – Harvard Apparatus GmbH, March-Hugstetten, Germany). The temporal sclera was punctured with a 30G needle just below the ora serrata. A 34G blunt end needle mounted on a 5 µL Hamilton syringe (Hamilton Bonaduz AG, Bonaduz, Switzerland), containing the AAV solution, was carefully inserted into the vitreous or the subretinal space through the puncture site using a micromanipulator (H. Saur Laborbedarf, Reutlingen, Germany). A cover slip was placed on top of the Lacrinorm gel for better visibility of the needle. Photoreceptors were transduced by a transvitreal subretinal injection of 1 µL AAV solution (1×10^9^ vg/uL) nasally to the optic nerve. Inner retinal neurons were transduced by an intravitreal injection of 1-4×10^9^ vg/μL AAVs (in 1 µL) above the optic nerve head. AAV solutions contained 10% fluorescein (0.1 mg/mL, Akorn Inc., IL, USA) to visually control the injections. After the procedure, the anesthesia was reversed by the subcutaneous injection of Atipamezol (1.67 mg/kg, Graeub, Bern, Switzerland) and mice were placed on a heating pad until fully awake. Wildtype mice were injected at 4-6 weeks of age and *Rho^P23H/+^* mice were injected at 4 weeks of age.

One to three weeks after injection, mice were subjected to fluorescent funduscopy and optical coherence tomography (OCT) using the Micron IV system (Phoenix Research Labs, Pleasanton, CA, USA) as described (*108*). Anesthesia and pupil dilation were performed as described above. Eyes with no transgene expression, injection-inflicted retinal damage or bleeding were excluded from the study.

### Immunofluorescence

Immunofluorescence was performed as previously described (*26*). Briefly, eyes were marked nasally, enucleated and fixed for 1.5 h in 4% paraformaldehyde (PFA) in PBS at 4°C. The cornea and lens were removed, and the eyes were fixed for another 2.5 h in PFA, followed by cryoprotection in 30% sucrose for 2 h, embedding and freezing in O.C.T. medium (O.C.T., 81-0771-00, Biosystems Switzerland AG, Nussloch, Germany). A Leica cryostat (Biosystems Switzerland AG, CM1860) was used to prepare 12 µm thick sections. Cryo-sections were blocked using PBS containing 3% normal goat serum (Sigma Aldrich, G9023, St. Louis, MO, USA) and 0.3% Triton X-100 (Sigma Aldrich, T9284) followed by incubation with the following primary antibodies overnight at 4°C: rabbit anti-ARR3 (1:1000; Merck Millipore Chemicals, Darmstadt, Germany; ab15282), mouse anti-CALB2 (1:1000; Merck Millipore Chemicals; MAB1568) and rabbit anti-RBPMS (1:200; Phosphosolutions, Aurora, CO, USA; 1830-RBPMS). After washing with PBS, the secondary antibodies (goat anti-rabbit IgG Alexa Fluor 568, A11011; Anti-mouse IgG Alexa Fluor 568, A11004; ThermoFisher (Waltham, MA, USA); 1:500) were applied for 1 h at room temperature. Mowiol (Mowiol 4-88, 475904, Calbiochem, Darmstadt, Germany) antifade medium was used to mount the sections. Fluorescence images were acquired with a confocal microscope (Andor BC43 CF, Oxford instruments, Abingdon, UK). The images were processed using ImageJ (Version 1.54f, National Institutes of Health).

### Morphology

For morphological analysis, eyes were fixed in 2.5% glutaraldehyde in 0.1 M cacodylate buffer, pH 7.2, at 4°C overnight. For each eye the posterior and inferior half were prepared, post-fixed in 1% osmium tetroxide in 0.1 M cacodylate buffer and embedded in EPON 812 (Biolyst Scientific, Morgantown, PA, USA) as described (*109*). Sections (500 nm) were cut through the optic nerve head, stained with toluidine blue and imaged by light microscopy (AxioImager Z2, Carl Zeiss AG, Feldbach, Switzerland).

### Acute eye section preparations for *ex vivo* imaging

The mice were housed in a room with a reversed 12/12 h light/dark cycle for at least 10 days before the experiments. Acute eye sections of wildtype mice were prepared at 8-12 weeks of age and of *Rho^P23H/+^*at 8 weeks of age. Mice were euthanized at 10 am during the subjective night. Eyes were marked nasally and embedded in 3% agarose (Type III-A, Sigma, A9793) dissolved in freshly prepared artificial cerebrospinal fluid (ACSF) containing: 126 mM NaCl, 3 mM KCl, 2 mM CaCl_2_, 1.25 mM NaH_2_PO_4_, 26 mM NaHCO_3_, 2 mM MgSO_4_, 6 mM D-glucose, bubbled with oxycarbon (95% O_2_ / 5% CO_2_, 800001855, Linde Gas Schweiz, Dagmersellen, Switzerland). Eyes were cut temporal to nasal into 300 µm thick acute sections in ice-cold ACSF on a microtome with a horizontally oscillating blade (Microm HM 650 V, Histocom AG, Zug, Switzerland) using the following settings: feed = 300, trim = 25, frequency = 85, amplitude = 1, speed = 7. The sections were transferred to a chamber containing ACSF that was continuously bubbled with oxycarbon and left to equilibrate for 2 h at RT. Ten to 15 min before imaging, the sections were transferred to the imaging chamber, to adapt to the imaging temperature while the ACSF was slowly warmed to 35°C. The section was held in position by a slice anchor (SS SLIC HD F/RC-26 CHAM 1.5 mm, 640251, MultiChannel Systems HEKA, Holliston, USA) to prevent movement during imaging. The imaging chamber (approximate volume: 2.5 mL) was perfused with continuously oxygenated ACSF at a flow rate of approximately 1.7 mL/min and the temperature was maintained at 35°C using a temperature controller (V TC05, Luigs & Neumann GmbH, Ratingen, Germany). Imaging was performed at the depth of 30-80 µm below the section surface.

### Two-photon excitation laser scanning microscopy (2P) and two-photon fluorescence lifetime microscopy (2P-FLIM)

A custom-made two-photon laser scanning microscope (*110*) equipped with a tunable femtosecond pulsed laser (Chameleon Discovery NX with Total Power Control, Coherent, Saxonburg, PA, USA) and a 20x water immersion objective (W Plan-Apochromat 20x/1.0, Zeiss) was used for image acquisition during two-photon excitation laser scanning microscopy (2P) and two-photon fluorescence lifetime microscopy (2P-FLIM). Excitation and emission beam paths were separated by a dichroic mirror (KS93 Cold Light Mirror, 685 nm LP, QiOptiq, Rhyl, UK). Dichroic mirrors at 506nm (F38-506; AHF Analysentechnik, Tübingen, Germany) and 560 nm (F38-560; AHF Analysentechnik) further separated the emission light into colored components, that were then focused on GaAsP photomultipliers (H10770PA-40sel; Hamamatsu Photonics, Hamamatsu, Japan) equipped with filters for cyan (Brightline HC 475/50; AHF Analysentechnik), green/yellow (Brightline HC 542/50; AHF Analysentechnik) and red (Brightline HC 607/70; AHF Analysentechnik) wavelengths. ScanImage 3.8 (*111*) and a custom-written LabVIEW software (Version 2012; National Instruments) were used for image control and data acquisition. For FLIM acquisition in a given channel, the GaAsP photomultiplier was replaced with a hybrid photomultiplier detector (PMA Hybrid 40 mod, PicoQuant, Berlin, Germany), and the signal collected and processed via a custom designed FLIM acquisition hardware and software based on an FPGA evaluation board (*112*). For FLIM in the cyan channel, an additional Brightline HC 475/50 filter was employed to eliminate any residual contribution from second harmonics generation.

### 2P Calcium imaging and analysis

Acute eye sections expressing GCaMP6s in rod photoreceptors and mCherry in RPE cells were co-excited with 940 nm and 1040 nm and emitted fluorescence intensity was acquired at a 0.74 Hz frame rate and 512 x 512 pixel resolution.

To measure Ca^2+^ levels in rod photoreceptors, GCaMP6s was excited at 940 nm with a laser power of 8-23 mW. Emitted fluorescence intensity was acquired at a 5.92 Hz frame rate and 128 x 128 pixel resolution. Rod photoreceptors were imaged for 10 s with 30 s recovery under continuous ACSF flow. To evaluate the contribution of calcium influx through cGMP-gated channels in photoreceptor cells, 2 mM 8-pCPT-cGMP (Table S1) was added to the ACSF and photoreceptors were continuously imaged for 200 s. For data analysis regions of interest (ROI) were manually outlined around photoreceptor segments using ImageJ. Further data analysis was conducted using a custom-made MATLAB (MathWorks, R2018b) script (https://gitlab.com/einlabzurich/dffanalysis). The Ca^2+^ traces (ΔF/F) were calculated from each ROI using the first frame of each experiment as baseline.

### 2P-FLIM of metabolic biosensors for ATP, lactate and NADH/NAD^+^

Imaging of the metabolic biosensors was performed using various pharmacological protocols (Table 1) at 2.96 Hz frame rate and 128 x 128 pixel resolution. The fluorescence lifetime of the sensors was collected at intervals of 25 s. 30 images (for the acquisition of fluorescence lifetime of LiLac and Peredox) or 20 images (for ATeam1.03) were averaged within the 25 s time interval. ATeam 1.03 and LiLac were excited at 870 nm using 3-12 mW laser power. Peredox was excited at 800 nm wavelength using 5 -13 mW laser power. Detailed information about imaging protocols and drugs are listed in Table 1 and Table S1, respectively.

### 2P-FLIM analysis

FLIM images were analyzed using custom-made MATLAB (MathWorks, R2020a) scripts based on the FLIMfit 5.1.1 library (*113*) (https://gitlab.com/einlabzurich/flimanalysis) using biexponential fitting and the method-of-moments (MOM), as previously described (*112*). Although all analyses were broadly consistent, we noticed the presence of very-short lived autofluorescence, especially in dimmer areas corresponding to the photoreceptor outer segments. To reduce its impact, regions-of-interests excluding outer segments were selected, and the average fluorescence decays in this region was analyzed. To analyze fluorescence lifetime changes in the inner retina, the average fluorescence decay of the whole frame was analyzed.

To further minimize the contribution of short-lived contaminations, we first detected the center-of-mass of the IRF (Instrument Response Function), and aligned the decays such that it would fall 250 ps from the start of the time-scale. We then excluded the first 750 ps of the signal (corresponding to roughly 500 ps after the peak of excitation), as they would be the most affected by the contaminating signal. Finally, we conducted a MOM analysis on the cropped decays (“MOM cropped”, which we will refer to as τ_MC_).

The slopes were calculated using the linear function: y = mx + b (where m represents the slope) in GraphPad Prism version 10.5.0 (GraphPad Software, San Diego, CA, USA).

### Statistical analysis and presentation

The numbers of mice (n) and slices (N) are detailed in the figure legends. GraphPad Prism Version 10.5.0 was used for data visualization, and the Mann-Whitney nonparametric test was used to compare data.

## Acknowledgements

We thank the Viral Vector Facility (VVF) of the Neuroscience Center Zurich (ZNZ) for the production of all AAVs and the Laboratory Animal Services Center (LASC) of the University of Zurich for animal care. We thank Luca Merolla for providing tissue sections for morphological analysis. We also thank Felipe Barros for scientific discussions.

## Author contributions

G.M.W., M.S., V.T., B.W. and C.G. conceived the study. G.M.W. performed AAV injections and 2P/2P-FLIM experiments. G.M.W., L.R. and R.M.M. analyzed and visualized the 2P/2P-FLIM data. G.M.W. performed and visualized immunofluorescence and morphological analysis. G.M.W and C.G. wrote the manuscript. All authors contributed to the project design and interpretation of results. All authors read and approved the final manuscript.

## Data availability

The code utilized to perform the analysis of the Ca^2+^ data is available at https://gitlab.com/einlabzurich/dffanalysis and the code utilized to perform FLIM analysis is available at https://gitlab.com/einlabzurich/flimanalysis.

Any additional information required to reanalyze the data reported in this paper are available from the corresponding author upon request.

## Supplementary Figures

**Fig. S1:** The flow-rate in the imaging chamber and the cell-type specificity of biosensor expression. **A,** Representative intensity trace of fluorescein. Baseline fluorescence intensity during inflow of ACSF was recorded for 3 min, followed by inflow of fluorescein (1×10^−5^ M fluorescein in ACSF; *green arrow*: start of inflow) for 10 min. Fluorescein arrived in the chamber after approx. 1.4 min of inflow (*blue arrow*). After 10 min, fluorescein was washed out with ACSF (*red arrow*: start of outwash). **B,** Subretinal delivery of AAV::mOP-ATeam1.03 (left) and retinal cross-sections showing ATeam1.03 fluorescence (green) and cone arrestin immunofluorescence (right, ARR3, red). White arrows indicate ARR3-positive but ATeam1.03-negative cells. **C,** Intravitreal delivery of AAV::hSyn1-ATeam1.03 (left) and cross-sections showing ATeam1.03 expression (green) and calretinin (CALB2, red, right upper panel) or RNA binding protein with multiple splicing (RBPMS, red, right lower panel) immunofluorescence. White arrows indicate cells expressing ATeam1.03 and the amacrine cell marker CALB2 or the ganglion cell marker RBPMS. Scale bars: 20 µm. GCL: ganglion cell layer. INL: inner nuclear layer. IPL: inner plexiform layer. ONL: outer nuclear layer. OPL: outer plexiform layer. PS: photoreceptor segments.

**Fig. S2:** Stability of ATeam1.03 fluorescence lifetime and recovery after oxidative phosphorylation block in rods of wildtype mice (129S6). **A**, Lifetime trace (mean ± SD; N = 3, n = 3) of ATeam1.03 in rods during the ‘control protocol’ (Table 1). **B,** Lifetime trace (mean ± SD; N = 3, n = 3) of ATeam1.03 in rods during the ‘NaN_3_ outwash protocol’ (Table 1).

**Fig. S3:** ATP dynamics in rods supplied with glutamine of wildtype mice (129S6). **A,** Schematic representation of glutamine metabolism. **B,** 2P-FLIM images (τ_MC_) of ATeam1.03 in rods at different time points during the ‘glutamine supplementation protocol’. Scale bar: 10 µm. **C,** Lifetime traces of ATeam1.03 in rods (N = 7, n = 3) during the ‘glutamine supplementation protocol’ (mean ± SD). *Black dashed line*: slope used to calculate ATP depletion rates. **D,** Comparison of the ATP depletion rates (Δτ_MC_/min) during the ‘lactate supplementation protocol’ (data from Fig. 2D’’’, *green*) and the ‘glutamine supplementation protocol’ (*light green*) in rods. Shown are individual values and means ± SD. Mann-Whitney nonparametric test.

**Fig. S4:** Alternative fuels for ATP production in inner retinal neurons of wildtype mice (129S6). **A,** Schematic representation of cellular metabolism. Monocarboxylate transporters (MCT) were blocked with AR-C155858 (AR-C), glucose (Glc) uptake with cytoB and OXPHOS with NaN3. **B,** 2P-FLIM images (τ_MC_) of ATeam1.03 in inner retinal neurons at different time points during the ‘lactate transport block protocol’ (Table 1). Scale bar: 10 µm. **C,** Lifetime trace of ATeam1.03 in inner retinal neurons during the lactate transport block protocol (mean ± SD; N = 10, n = 5). Black dashed line: slope used to calculate ATP depletion rates. **D,** Comparison of ATP depletion rates (Δτ_MC_/min) during the ‘glucose depletion protocol’ (data from Fig. 2E’’’, dark blue) and the ‘lactate transport block protocol’ (turquoise). Shown are individual values and means ± SD. Mann-Whitney nonparametric test. **E,** Schematic representation of glycogen metabolism and the inhibition of glycogen phosphorylase with 1,4-Dideoxy-1,4-imino-D-arabinitol hydrochloride (DAB) to prevent the breakdown of glycogen to enter glycolysis (Table 1). **F,** 2P-FLIM images (τ_MC_) of ATeam1.03 in inner retinal neurons at different time points during the ‘glycogen phosphorylase block protocol’ (Table 1). Scale bar: 10 µm. **G,** Lifetime trace of ATeam1.03 in inner retinal neurons during the ‘glycogen phosphorylase block protocol’ (mean ± SD; N = 8, n = 5). Black dashed line: slope used to calculate ATP depletion rates. **H,** Comparison of ATP depletion rates (Δτ_MC_/min) during the ‘glucose depletion protocol’ (data from Fig. 2E’’’, dark blue) and the ‘glycogen phosphorylase block protocol’ (light blue). Shown are individual values and means ± SD. Mann-Whitney nonparametric test.

**Fig. S5:** Redox state measurements in rods and neurons of the inner retina of wildtype mice (129S6). **A,** Schematic representation of the NADH/NAD^+^ sensor Peredox (adapted from Hung and Yellen (54)). The lifetime (τ) of Peredox increases when the NADH/NAD^+^ ratio is high (glycolytic) and decreases when the NADH/NAD^+^ ratio is low (oxidative). **B,** Schematic representation of main mechanisms influencing the cytosolic NADH/NAD^+^ ratio in cells. **C,** 2P-FLIM images (τ_MC_) of Peredox in rods (upper panel) and inner retinal neurons (lower panel) at baseline. Scale bars: 10 µm. **D,** Comparison of the lifetime (τ_MC_) (Δτ_MC_/min) of Peredox at baseline levels in 6mM glucose in rods (*green*) and inner retinal neurons (*blue*). Shown are individual values and means ± SD. Statistics: Mann-Whitney nonparametric test.

**Fig. S6:** Lactate production and consumption by blocking different MCTs in rods of wildtype mice (129S6). A, Schematic representation of cellular metabolism, illustrating lactate transport via MCTs and their blocking with AR-C155858 (MCT1/2) and Syrosingopine (MCT1/MCT4). B, 2P-FLIM images (τ_MC_) of LiLac in rods at different time points during the ‘lactate production/consumption protocol 2’ (Table 1). Scale bar: 10 µm. C, Lifetime traces (means ± SD) of LiLac in rods (N = 7; n = 3) during the ‘lactate production/consumption protocol 2’. *Black dashed lines*: slope used to calculate the changes in the lifetime in LiLac shown in panels D, E. D, E, Lifetime changes (Δτ_MC_/min) in LiLac during the ‘lactate consumption/production protocol 1’ while blocking MCT1/2 (data from Fig. 4 E, F, *green*) and during the ‘lactate consumption/production protocol 2’ while blocking MCT1, 2 and 4 (*bright green*), in the presence of glucose (D; slope 1; lactate production) and in the absence of glucose (E; slope 2; lactate consumption). F, Lifetime traces (means ± SD) of LiLac in rods (N = 2; n = 1) during the ‘Syrosingopine test protocol’. *Black dashed lines*: slope used to calculate the changes in the lifetime in LiLac shown in panels G,H. G, H, Lifetime changes (Δτ_MC_/min) in LiLac during the ‘lactate consumption/production protocol 1’ while blocking MCT1/2 (data from Fig. 4 E, F, *green*) and during the ‘lactate consumption/production protocol 2’ while blocking MCT1, 2 and 4 (*light green*), in the presence of glucose (G; slope 1; lactate production) and in the absence of glucose (H; slope 2; lactate consumption). Shown are individual values and means ± SD. Mann-Whitney nonparametric test.

**Fig. S7:** Light induced response of rods in *Rho^P23H/+^* mice during 2P. **A,** Representative images of the retinal morphology of wildtype (WT, C57B6/J) and *Rho^P23H/+^* mice at 8 weeks of age. **B,** Schematic representation of the experimental timeline. Mice received a subretinal injection at 4 weeks of age. 2P of acute eye sections was performed at 8 weeks of age. **C,** Representative 2P images of acute eye sections expressing GCaMP6s in rods of *Rho^P23H/+^* mice. Left: maximum intensity at 0 s. Middle and right: intensity weighted images at 0 s (middle) and 10 s (right) of the imaging series. Red boxes mark the analyzed area (photoreceptor segments). **D,** GCaMP6s traces (mean ± SD) imaged in photoreceptor segments of *Rho^P23H/+^* mice (N = 9, n = 4) during 10 s of 2P imaging followed by 30 s recovery. **E,** GCaMP6s trace (mean ± SD) of the photoreceptor segments of *Rho^P23H/+^* mice (N = 4, n = 3) in the presence of 2 mM 8-pCPT-cGMP. Scale bars: 10 μm. ONL: outer nuclear layer. OPL: outer plexiform layer. PS: photoreceptor segments.

## References

1. A. Ames, Energy requirements of CNS cells as related to their function and to their vulnerability to ischemia: a commentary based on studies on retina. Can J Physiol Pharmacol 70 **Suppl**, S158–164 (1992).

2. M. T. Wong-Riley, Energy metabolism of the visual system. Eye Brain 2, 99–116 (2010).

3. H. Liu, V. Prokosch, Energy Metabolism in the Inner Retina in Health and Glaucoma. International Journal of Molecular Sciences 22, 3689 (2021).

4. H. Okawa, A. P. Sampath, S. B. Laughlin, G. L. Fain, ATP consumption by mammalian rod photoreceptors in darkness and in light. Curr Biol 18, 1917–1921 (2008).

5. N. T. Ingram, G. L. Fain, A. P. Sampath, Elevated energy requirement of cone photoreceptors. Proc Natl Acad Sci U S A 117, 19599–19603 (2020).

6. J. D. Linton, L. C. Holzhausen, N. Babai, H. Song, K. J. Miyagishima, G. W. Stearns, K. Lindsay, J. Wei, A. O. Chertov, T. A. Peters, R. Caffe, H. Pluk, M. W. Seeliger, N. Tanimoto, K. Fong, L. Bolton, D. L. Kuok, I. R. Sweet, T. M. Bartoletti, R. A. Radu, G. H. Travis, W. N. Zagotta, E. Townes-Anderson, E. Parker, C. E. Van der Zee, A. P. Sampath, M. Sokolov, W. B. Thoreson, J. B. Hurley, Flow of energy in the outer retina in darkness and in light. Proc Natl Acad Sci U S A 107, 8599–8604 (2010).

7. J. Hanna, L. A. David, Y. Touahri, T. Fleming, R. A. Screaton, C. Schuurmans, Beyond Genetics: The Role of Metabolism in Photoreceptor Survival, Development and Repair. Front Cell Dev Biol 10, 887764 (2022).

8. C. J. Medrano, D. A. Fox, Oxygen consumption in the rat outer and inner retina: light- and pharmacologically-induced inhibition. Exp Eye Res 61, 273–284 (1995).

9. D.-Y. Yu, S. J. Cringle, Retinal degeneration and local oxygen metabolism. Experimental Eye Research 80, 745–751 (2005).

10. O. Warburg, Über den Stoffwechsel der Carcinomzelle. Naturwissenschaften 12, 1131–1137 (1924).

11. B. S. Winkler, Glycolytic and oxidative metabolism in relation to retinal function. Journal of General Physiology 77, 667–692 (1981).

12. J. B. Hurley, Retina Metabolism and Metabolism in the Pigmented Epithelium: A Busy Intersection. Annu Rev Vis Sci 7, 665–692 (2021).

13. Y. Chinchore, T. Begaj, D. Wu, E. Drokhlyansky, C. L. Cepko, Glycolytic reliance promotes anabolism in photoreceptors. Elife 6, (2017).

14. A. Rajala, Y. Wang, R. S. Brush, K. Tsantilas, C. S. R. Jankowski, K. J. Lindsay, J. D. Linton, J. B. Hurley, R. E. Anderson, R. V. S. Rajala, Pyruvate kinase M2 regulates photoreceptor structure, function, and viability. Cell Death & Disease 9, 240 (2018).

15. T. J. Wubben, M. Pawar, A. Smith, K. Toolan, H. Hager, C. G. Besirli, Photoreceptor metabolic reprogramming provides survival advantage in acute stress while causing chronic degeneration. Scientific Reports 7, 17863 (2017).

16. A. Rajala, Y. Wang, K. Soni, R. V. S. Rajala, Pyruvate kinase M2 isoform deletion in cone photoreceptors results in age-related cone degeneration. Cell Death & Disease 9, 737 (2018).

17. A. Rajala, M. A. Bhat, K. Teel, G. K. Gopinadhan Nair, L. Purcell, R. V. S. Rajala, The function of lactate dehydrogenase A in retinal neurons: implications to retinal degenerative diseases. PNAS Nexus 2, (2023).

18. R. V. S. Rajala, A. Rajala, C. Kooker, Y. Wang, R. E. Anderson, The Warburg Effect Mediator Pyruvate Kinase M2 Expression and Regulation in the Retina. Scientific Reports 6, 37727 (2016).

19. K. J. Lindsay, J. Du, S. R. Sloat, L. Contreras, J. D. Linton, S. J. Turner, M. Sadilek, J. Satrústegui, J. B. Hurley, Pyruvate kinase and aspartate-glutamate carrier distributions reveal key metabolic links between neurons and glia in retina. Proc Natl Acad Sci U S A 111, 15579–15584 (2014).

20. L. Petit, S. Ma, J. Cipi, S. Y. Cheng, M. Zieger, N. Hay, C. Punzo, Aerobic Glycolysis Is Essential for Normal Rod Function and Controls Secondary Cone Death in Retinitis Pigmentosa. Cell Rep 23, 2629–2642 (2018).

21. R. J. Casson, J. P. M. Wood, G. Han, T. Kittipassorn, D. J. Peet, G. Chidlow, M-Type Pyruvate Kinase Isoforms and Lactate Dehydrogenase A in the Mammalian Retina: Metabolic Implications. Investigative Ophthalmology & Visual Science 57, 66–80 (2016).

22. R. Zhang, W. Shen, J. Du, M. C. Gillies, Selective knockdown of hexokinase 2 in rods leads to age-related photoreceptor degeneration and retinal metabolic remodeling. Cell Death Dis 11, 885 (2020).

23. E. M. Rueda, J. E. Johnson, Jr., A. Giddabasappa, A. Swaroop, M. J. Brooks, I. Sigel, S. Y. Chaney, D. A. Fox, The cellular and compartmental profile of mouse retinal glycolysis, tricarboxylic acid cycle, oxidative phosphorylation, and ∼P transferring kinases. Mol Vis 22, 847–885 (2016).

24. M. A. Kanow, M. M. Giarmarco, C. S. Jankowski, K. Tsantilas, A. L. Engel, J. Du, J. D. Linton, C. C. Farnsworth, S. R. Sloat, A. Rountree, I. R. Sweet, K. J. Lindsay, E. D. Parker, S. E. Brockerhoff, M. Sadilek, J. R. Chao, J. B. Hurley, Biochemical adaptations of the retina and retinal pigment epithelium support a metabolic ecosystem in the vertebrate eye. Elife 6, (2017).

25. J. Du, W. Cleghorn, L. Contreras, J. D. Linton, G. C. Chan, A. O. Chertov, T. Saheki, V. Govindaraju, M. Sadilek, J. Satrústegui, J. B. Hurley, Cytosolic reducing power preserves glutamate in retina. Proc Natl Acad Sci U S A 110, 18501–18506 (2013).

26. V. Todorova, M. F. Stauffacher, L. Ravotto, S. Nötzli, D. Karademir, L. J. A. Ebner, C. Imsand, L. Merolla, S. M. Hauck, M. Samardzija, A. S. Saab, L. F. Barros, B. Weber, C. Grimm, Deficits in mitochondrial TCA cycle and OXPHOS precede rod photoreceptor degeneration during chronic HIF activation. Molecular Neurodegeneration 18, 15 (2023).

27. V. Calbiague-Garcia, Y. Chen, B. Cádiz, F. Tapia, F. Paquet-Durand, O. Schmachtenberg, Extracellular lactate as an alternative energy source for retinal bipolar cells. J Biol Chem 300, 106794 (2024).

28. V. Calbiague García, Y. Chen, B. Cádiz, L. Wang, F. Paquet-Durand, O. Schmachtenberg, Imaging of lactate metabolism in retinal Müller cells with a FRET nanosensor. Exp Eye Res 226, 109352 (2023).

29. L. C. Chandler, A. Gardner, C. L. Cepko, RPE-specific MCT2 expression promotes cone survival in models of retinitis pigmentosa. Proceedings of the National Academy of Sciences 122, e2421978122 (2025).

30. S. Y. Lunt, M. G. Vander Heiden, Aerobic glycolysis: meeting the metabolic requirements of cell proliferation. Annu Rev Cell Dev Biol 27, 441–464 (2011).

31. L. L. Daniele, J. Y. S. Han, I. S. Samuels, R. Komirisetty, N. Mehta, J. L. McCord, M. Yu, Y. Wang, K. Boesze-Battaglia, B. A. Bell, J. Du, N. S. Peachey, N. J. Philp, Glucose uptake by GLUT1 in photoreceptors is essential for outer segment renewal and rod photoreceptor survival. The FASEB Journal 36, e22428 (2022).

32. A. Swarup, I. S. Samuels, B. A. Bell, J. Y. S. Han, J. Du, E. Massenzio, E. D. Abel, K. Boesze-Battaglia, N. S. Peachey, N. J. Philp, Modulating GLUT1 expression in retinal pigment epithelium decreases glucose levels in the retina: impact on photoreceptors and Müller glial cells. Am J Physiol Cell Physiol 316, C121–c133 (2019).

33. L. Wang, P. Tornquist, A. Bill, Glucose metabolism in pig outer retina in light and darkness. Acta Physiol Scand 160, 75–81 (1997).

34. L. Wang, M. Kondo, A. Bill, Glucose metabolism in cat outer retina. Effects of light and hyperoxia. Invest Ophthalmol Vis Sci 38, 48–55 (1997).

35. F. O. Viegas, S. C. F. Neuhauss, A Metabolic Landscape for Maintaining Retina Integrity and Function. Front Mol Neurosci 14, 656000 (2021).

36. Richard H. Masland, The Neuronal Organization of the Retina. Neuron 76, 266–280 (2012).

37. H. Wässle, Parallel processing in the mammalian retina. Nature Reviews Neuroscience 5, 747–757 (2004).

38. L. Wang, P. Törnquist, A. Bill, Glucose metabolism of the inner retina in pigs in darkness and light. Acta Physiol Scand 160, 71–74 (1997).

39. R. J. Casson, G. Chidlow, J. G. Crowston, P. A. Williams, J. P. M. Wood, Retinal energy metabolism in health and glaucoma. Progress in Retinal and Eye Research 81, 100881 (2021).

40. W. W. Pan, T. J. Wubben, C. G. Besirli, Photoreceptor metabolic reprogramming: current understanding and therapeutic implications. Communications Biology 4, 245 (2021).

41. D. M. Inman, M. Harun-Or-Rashid, Metabolic Vulnerability in the Neurodegenerative Disease Glaucoma. Front Neurosci 11, 146 (2017).

42. Y. Liu, X. Wang, R. Gong, G. Xu, M. Zhu, Overexpression of Rhodopsin or Its Mutants Leads to Energy Metabolism Dysfunction in 661w Cells. Invest Ophthalmol Vis Sci 63, 2 (2022).

43. Y. Tomita, C. Qiu, E. Bull, W. Allen, Y. Kotoda, S. Talukdar, L. E. H. Smith, Z. Fu, Müller glial responses compensate for degenerating photoreceptors in retinitis pigmentosa. Exp Mol Med 53, 1748–1758 (2021).

44. T. McLaughlin, A. Medina, J. Perkins, M. Yera, J. J. Wang, S. X. Zhang, Cellular stress signaling and the unfolded protein response in retinal degeneration: mechanisms and therapeutic implications. Molecular Neurodegeneration 17, 25 (2022).

45. N. D. Nolan, S. M. Caruso, X. Cui, S. H. Tsang, Renormalization of metabolic coupling treats age-related degenerative disorders: an oxidative RPE niche fuels the more glycolytic photoreceptors. Eye 36, 278–283 (2022).

46. R. Datta, T. M. Heaster, J. T. Sharick, A. A. Gillette, M. C. Skala, Fluorescence lifetime imaging microscopy: fundamentals and advances in instrumentation, analysis, and applications. J Biomed Opt 25, 1–43 (2020).

47. G. Yellen, R. Mongeon, Quantitative two-photon imaging of fluorescent biosensors. Curr Opin Chem Biol 27, 24–30 (2015).

48. M. Hoon, H. Okawa, L. Della Santina, R. O. Wong, Functional architecture of the retina: development and disease. Prog Retin Eye Res 42, 44–84 (2014).

49. N. Esumi, Y. Oshima, Y. Li, P. A. Campochiaro, D. J. Zack, Analysis of the VMD2 promoter and implication of E-box binding factors in its regulation. J Biol Chem 279, 19064–19073 (2004).

50. W. A. Beltran, S. L. Boye, S. E. Boye, V. A. Chiodo, A. S. Lewin, W. W. Hauswirth, G. D. Aguirre, rAAV2/5 gene-targeting to rods:dose-dependent efficiency and complications associated with different promoters. Gene Ther 17, 1162–1174 (2010).

51. T. W. Chen, T. J. Wardill, Y. Sun, S. R. Pulver, S. L. Renninger, A. Baohan, E. R. Schreiter, R. A. Kerr, M. B. Orger, V. Jayaraman, L. L. Looger, K. Svoboda, D. S. Kim, Ultrasensitive fluorescent proteins for imaging neuronal activity. Nature 499, 295–300 (2013).

52. L. Stryer, Visual excitation and recovery. Journal of Biological Chemistry 266, 10711–10714 (1991).

53. P. Zhang, M. Goswami, R. J. Zawadzki, E. N. Pugh, Jr., The Photosensitivity of Rhodopsin Bleaching and Light-Induced Increases of Fundus Reflectance in Mice Measured In Vivo With Scanning Laser Ophthalmoscopy. Invest Ophthalmol Vis Sci 57, 3650–3664 (2016).

54. S. Yokoyama, R. Yokoyama, Chapter 6 Comparative molecular biology of visual pigments (Handbook of Biological Physics, North-Holland, 2000), vol. 3, pp. 257–296.

55. T. Euler, K. Franke, T. Baden, Studying a Light Sensor with Light: Multiphoton Imaging in the Retina (Multiphoton Microscopy, Springer New York, New York, NY, 2019), pp. 225–250.

56. W. Denk, P. B. Detwiler, Optical recording of light-evoked calcium signals in the functionally intact retina. Proc Natl Acad Sci U S A 96, 7035–7040 (1999).

57. G. Palczewska, F. Vinberg, P. Stremplewski, M. P. Bircher, D. Salom, K. Komar, J. Zhang, M. Cascella, M. Wojtkowski, V. J. Kefalov, K. Palczewski, Human infrared vision is triggered by two-photon chromophore isomerization. Proceedings of the National Academy of Sciences 111, E5445–E5454 (2014).

58. P. Pliushcheuskaya, S. Kesh, E. Kaufmann, S. Wucherpfennig, F. Schwede, G. Künze, V. Nache, Similar Binding Modes of cGMP Analogues Limit Selectivity in Modulating Retinal CNG Channels via the Cyclic Nucleotide-Binding Domain. ACS Chemical Neuroscience 15, 1652–1668 (2024).

59. J.-Y. Wei, E. D. Cohen, H.-G. Genieser, C. J. Barnstable, Substituted cGMP analogs can act as selective agonists of the rod photoreceptor cGMP-gated cation channel. Journal of Molecular Neuroscience 10, 53–64 (1998).

60. H. Imamura, K. P. Nhat, H. Togawa, K. Saito, R. Iino, Y. Kato-Yamada, T. Nagai, H. Noji, Visualization of ATP levels inside single living cells with fluorescence resonance energy transfer-based genetically encoded indicators. Proc Natl Acad Sci U S A 106, 15651–15656 (2009).

61. K. Bogucka, L. Wojtczak, Effect of sodium azide on oxidation and phosphorylation processes in rat-liver mitochondria. Biochimica et Biophysica Acta (BBA) - Enzymology and Biological Oxidation 122, 381–392 (1966).

62. F. Baeza-Lehnert, A. S. Saab, R. Gutiérrez, V. Larenas, E. Díaz, M. Horn, M. Vargas, L. Hösli, J. Stobart, J. Hirrlinger, B. Weber, L. F. Barros, Non-Canonical Control of Neuronal Energy Status by the Na(+) Pump. Cell Metab 29, 668–680.e664 (2019).

63. C. X. Bittner, R. Valdebenito, I. Ruminot, A. Loaiza, V. Larenas, T. Sotelo-Hitschfeld, H. Moldenhauer, A. San Martín, R. Gutiérrez, M. Zambrano, L. F. Barros, Fast and reversible stimulation of astrocytic glycolysis by K+ and a delayed and persistent effect of glutamate. J Neurosci 31, 4709–4713 (2011).

64. C. X. Bittner, A. Loaiza, I. Ruminot, V. Larenas, T. Sotelo-Hitschfeld, R. Gutiérrez, A. Córdova, R. Valdebenito, W. B. Frommer, L. F. Barros, High resolution measurement of the glycolytic rate. Front Neuroenergetics 2, (2010).

65. E. Y.-C. Kang, Y.-J. Tseng, W.-H. Peng, H.-C. Hung, P.-H. Lin, K. P. Montales, E. Sherman, J. Peregrin, E. H. Wang, C. Kang, Y.-C. Teng, C.-Y. Huang, C.-L. Tsai, I. Y.-F. Chang, J. Chen, G. Tezel, Y. He, T.-D. Li, L. Stiles, O. Shirihai, S. H. Tsang, C.-C. Lai, C.-N. Tsai, C.-S. Lin, N.-K. Wang, Disrupted energy metabolism is associated with retinal ganglion cell degeneration in autosomal dominant optic atrophy. Science Advances 12, eadx7815 (2026).

66. M. J. Ovens, A. J. Davies, M. C. Wilson, C. M. Murray, A. P. Halestrap, AR-C155858 is a potent inhibitor of monocarboxylate transporters MCT1 and MCT2 that binds to an intracellular site involving transmembrane helices 7-10. Biochem J 425, 523–530 (2010).

67. A. B. Walls, H. M. Sickmann, A. Brown, S. D. Bouman, B. Ransom, A. Schousboe, H. S. Waagepetersen, Characterization of 1,4-dideoxy-1,4-imino-d-arabinitol (DAB) as an inhibitor of brain glycogen shunt activity. J Neurochem 105, 1462–1470 (2008).

68. Y. P. Hung, J. G. Albeck, M. Tantama, G. Yellen, Imaging cytosolic NADH-NAD(+) redox state with a genetically encoded fluorescent biosensor. Cell Metab 14, 545–554 (2011).

69. R. Mongeon, V. Venkatachalam, G. Yellen, Cytosolic NADH-NAD(+) Redox Visualized in Brain Slices by Two-Photon Fluorescence Lifetime Biosensor Imaging. Antioxid Redox Signal 25, 553–563 (2016).

70. M. A. Walker, R. Tian, NAD(H) in mitochondrial energy transduction: implications for health and disease. Curr Opin Physiol 3, 101–109 (2018).

71. D. Koveal, P. C. Rosen, D. J. Meyer, C. M. Díaz-García, Y. Wang, L.-H. Cai, P. J. Chou, D. A. Weitz, G. Yellen, A high-throughput multiparameter screen for accelerated development and optimization of soluble genetically encoded fluorescent biosensors. Nature Communications 13, 2919 (2022).

72. C. M. Bisbach, D. T. Hass, E. D. Thomas, T. J. Cherry, J. B. Hurley, Monocarboxylate Transporter 1 (MCT1) Mediates Succinate Export in the Retina. Invest Ophthalmol Vis Sci 63, 1 (2022).

73. D. Benjamin, D. Robay, S. K. Hindupur, J. Pohlmann, M. Colombi, M. Y. El-Shemerly, S. M. Maira, C. Moroni, H. A. Lane, M. N. Hall, Dual Inhibition of the Lactate Transporters MCT1 and MCT4 Is Synthetic Lethal with Metformin due to NAD+ Depletion in Cancer Cells. Cell Rep 25, 3047–3058.e3044 (2018).

74. S. Sakami, T. Maeda, G. Bereta, K. Okano, M. Golczak, A. Sumaroka, A. J. Roman, A. V. Cideciyan, S. G. Jacobson, K. Palczewski, Probing mechanisms of photoreceptor degeneration in a new mouse model of the common form of autosomal dominant retinitis pigmentosa due to P23H opsin mutations. J Biol Chem 286, 10551–10567 (2011).

75. M. Aguilà, J. Bellingham, D. Athanasiou, D. Bevilacqua, Y. Duran, R. Maswood, D. A. Parfitt, T. Iwawaki, G. Spyrou, A. J. Smith, R. R. Ali, M. E. Cheetham, AAV-mediated ERdj5 overexpression protects against P23H rhodopsin toxicity. Hum Mol Genet 29, 1310–1318 (2020).

76. E.-J. Lee, P. Chan, L. Chea, K. Kim, R. J. Kaufman, J. H. Lin, ATF6 is required for efficient rhodopsin clearance and retinal homeostasis in the P23H rho retinitis pigmentosa mouse model. Scientific Reports 11, 16356 (2021).

77. W. Wang, A. Kini, Y. Wang, T. Liu, Y. Chen, E. Vukmanic, D. Emery, Y. Liu, X. Lu, L. Jin, S. J. Lee, P. Scott, X. Liu, K. Dean, Q. Lu, E. Fortuny, R. James, H. J. Kaplan, J. Du, D. C. Dean, Metabolic Deregulation of the Blood-Outer Retinal Barrier in Retinitis Pigmentosa. Cell Rep 28, 1323–1334.e1324 (2019).

78. M. G. Vander Heiden, L. C. Cantley, C. B. Thompson, Understanding the Warburg effect: the metabolic requirements of cell proliferation. Science 324, 1029–1033 (2009).

79. R. W. Young, The renewal of photoreceptor cell outer segments. J Cell Biol 33, 61–72 (1967).

80. C. M. Bisbach, D. T. Hass, B. M. Robbings, A. M. Rountree, M. Sadilek, I. R. Sweet, J. B. Hurley, Succinate Can Shuttle Reducing Power from the Hypoxic Retina to the O2-Rich Pigment Epithelium. Cell Reports 31, 107606 (2020).

81. P. Altea-Manzano, A. Vandekeere, J. Edwards-Hicks, M. Roldan, E. Abraham, X. Lleshi, A. N. Guerrieri, D. Berardi, J. Wills, J. M. Junior, A. Pantazi, J. C. Acosta, R. M. Sanchez-Martin, S.-M. Fendt, M. Martin-Hernandez, A. J. Finch, Reversal of mitochondrial malate dehydrogenase 2 enables anaplerosis via redox rescue in respiration-deficient cells. Molecular Cell 82, 4537–4547.e4537 (2022).

82. A. O. Chertov, L. Holzhausen, I. T. Kuok, D. Couron, E. Parker, J. D. Linton, M. Sadilek, I. R. Sweet, J. B. Hurley, Roles of Glucose in Photoreceptor Survival. Journal of Biological Chemistry 286, 34700–34711 (2011).

83. L. Pellerin, P. J. Magistretti, Glutamate uptake into astrocytes stimulates aerobic glycolysis: a mechanism coupling neuronal activity to glucose utilization. Proc Natl Acad Sci U S A 91, 10625–10629 (1994).

84. P. Mächler, M. T. Wyss, M. Elsayed, J. Stobart, R. Gutierrez, A. von Faber-Castell, V. Kaelin, M. Zuend, A. San Martín, I. Romero-Gómez, F. Baeza-Lehnert, S. Lengacher, B. L. Schneider, P. Aebischer, P. J. Magistretti, L. F. Barros, B. Weber, In Vivo Evidence for a Lactate Gradient from Astrocytes to Neurons. Cell Metab 23, 94–102 (2016).

85. J. B. Hurley, K. J. Lindsay, J. Du, Glucose, lactate, and shuttling of metabolites in vertebrate retinas. J Neurosci Res 93, 1079–1092 (2015).

86. S. Kügler, E. Kilic, M. Bähr, Human synapsin 1 gene promoter confers highly neuron-specific long-term transgene expression from an adenoviral vector in the adult rat brain depending on the transduced area. Gene Therapy 10, 337–347 (2003).

87. D.-Y. Yu, S. J. Cringle, C. Balaratnasingam, W. H. Morgan, P. K. Yu, E.-N. Su, Retinal ganglion cells: Energetics, compartmentation, axonal transport, cytoskeletons and vulnerability. Progress in Retinal and Eye Research 36, 217–246 (2013).

88. J. T. Ortega, T. Parmar, M. Carmena-Bargueño, H. Pérez-Sánchez, B. Jastrzebska, Flavonoids improve the stability and function of P23H rhodopsin slowing down the progression of retinitis pigmentosa in mice. J Neurosci Res 100, 1063–1083 (2022).

89. M. Azam, B. Jastrzebska, Mechanisms of Rhodopsin-Related Inherited Retinal Degeneration and Pharmacological Treatment Strategies. Cells 14, (2025).

90. C. Punzo, W. Xiong, C. L. Cepko, Loss of Daylight Vision in Retinal Degeneration: Are Oxidative Stress and Metabolic Dysregulation to Blame?*. Journal of Biological Chemistry 287, 1642–1648 (2012).

91. K. Komeima, B. S. Rogers, P. A. Campochiaro, Antioxidants slow photoreceptor cell death in mouse models of retinitis pigmentosa. J Cell Physiol 213, 809–815 (2007).

92. M. L. Acosta, Y. S. Shin, S. Ready, E. L. Fletcher, D. L. Christie, M. Kalloniatis, Retinal metabolic state of the proline-23-histidine rat model of retinitis pigmentosa. Am J Physiol Cell Physiol 298, C764–774 (2010).

93. Y. Lin, Y. Wang, P. F. Li, Mutual regulation of lactate dehydrogenase and redox robustness. Front Physiol 13, 1038421 (2022).

94. W. Wang, S. J. Lee, P. A. Scott, X. Lu, D. Emery, Y. Liu, T. Ezashi, M. R. Roberts, J. W. Ross, H. J. Kaplan, D. C. Dean, Two-Step Reactivation of Dormant Cones in Retinitis Pigmentosa. Cell Rep 15, 372–385 (2016).

95. C. Punzo, K. Kornacker, C. L. Cepko, Stimulation of the insulin/mTOR pathway delays cone death in a mouse model of retinitis pigmentosa. Nature Neuroscience 12, 44–52 (2009).

96. N. Aït-Ali, R. Fridlich, G. Millet-Puel, E. Clérin, F. Delalande, C. Jaillard, F. Blond, L. Perrocheau, S. Reichman, L. C. Byrne, A. Olivier-Bandini, J. Bellalou, E. Moyse, F. Bouillaud, X. Nicol, D. Dalkara, A. van Dorsselaer, J. A. Sahel, T. Léveillard, Rod-derived cone viability factor promotes cone survival by stimulating aerobic glycolysis. Cell 161, 817–832 (2015).

97. L. Zhang, J. Du, S. Justus, C. W. Hsu, L. Bonet-Ponce, W. H. Wu, Y. T. Tsai, W. P. Wu, Y. Jia, J. K. Duong, V. B. Mahajan, C. S. Lin, S. Wang, J. B. Hurley, S. H. Tsang, Reprogramming metabolism by targeting sirtuin 6 attenuates retinal degeneration. J Clin Invest 126, 4659–4673 (2016).

98. Y. Chen, Y. Dong, J. Yan, L. Wang, S. Yu, K. Jiao, F. Paquet-Durand, Single-Cell Transcriptomic Profiling in Inherited Retinal Degeneration Reveals Distinct Metabolic Pathways in Rod and Cone Photoreceptors. International Journal of Molecular Sciences 23, 12170 (2022).

99. S. J. Lee, D. Emery, E. Vukmanic, Y. Wang, X. Lu, W. Wang, E. Fortuny, R. James, H. J. Kaplan, Y. Liu, J. Du, D. C. Dean, Metabolic transcriptomics dictate responses of cone photoreceptors to retinitis pigmentosa. Cell Rep 42, 113054 (2023).

100. D. T. Hass, C. M. Bisbach, M. Sadilek, I. R. Sweet, J. B. Hurley, Aerobic Glycolysis in Photoreceptors Supports Energy Demand in the Absence of Mitochondrial Coupling (Retinal Degenerative Diseases XIX, Springer International Publishing, Cham, 2023), pp. 435–441.

101. D. T. Hass, E. Giering, J. Y. S. Han, C. M. Bisbach, K. Pandey, B. M. Robbings, T. O. Mundinger, N. D. Nolan, S. H. Tsang, N. S. Peachey, N. J. Philp, J. B. Hurley, In vivo exchange of glucose and lactate between photoreceptors and the retinal pigment epithelium. eLife, (2025).

102. R. A. Linsenmeier, Effects of light and darkness on oxygen distribution and consumption in the cat retina. J Gen Physiol 88, 521–542 (1986).

103. J. L. Stobart, K. D. Ferrari, M. J. P. Barrett, M. J. Stobart, Z. J. Looser, A. S. Saab, B. Weber, Long-term In Vivo Calcium Imaging of Astrocytes Reveals Distinct Cellular Compartment Responses to Sensory Stimulation. Cerebral Cortex 28, 184–198 (2016).

104. J. Dernic, A. Eleftheriou, L. Vasilikos, M. Rauch, P. Imseng, H. S. Zanker, Z. J. Looser, R. M. Meister, F. Velasquez Moros, T. Kagan, T. Laviv, J.-C. Paterna, M. Arand, A. S. Saab, B. Weber, L. Ravotto, Abundance-biased codon diversification prevents recombination in AAV production and ensures robust in vivo expression of functional FRET sensors. Communications Biology 8, 1244 (2025).

105. C. N. Kay, R. C. Ryals, G. V. Aslanidi, S. H. Min, Q. Ruan, J. Sun, F. M. Dyka, D. Kasuga, A. E. Ayala, K. Van Vliet, M. Agbandje-McKenna, W. W. Hauswirth, S. L. Boye, S. E. Boye, Targeting photoreceptors via intravitreal delivery using novel, capsid-mutated AAV vectors. PLoS One 8, e62097 (2013).

106. C. A. Reid, K. J. Ertel, D. M. Lipinski, Improvement of Photoreceptor Targeting via Intravitreal Delivery in Mouse and Human Retina Using Combinatory rAAV2 Capsid Mutant Vectors. Invest Ophthalmol Vis Sci 58, 6429–6439 (2017).

107. S. J. Gasparini, K. Tessmer, M. Reh, S. Wieneke, M. Carido, M. Völkner, O. Borsch, A. Swiersy, M. Zuzic, O. Goureau, T. Kurth, V. Busskamp, G. Zeck, M. O. Karl, M. Ader, Transplanted human cones incorporate into the retina and function in a murine cone degeneration model. J Clin Invest 132, (2022).

108. P. Geiger, M. Barben, C. Grimm, M. Samardzija, Blue light-induced retinal lesions, intraretinal vascular leakage and edema formation in the all-cone mouse retina. Cell Death & Disease 6, e1985–e1985 (2015).

109. M. Samardzija, A. Wenzel, S. Aufenberg, M. Thiersch, C. Remé, C. Grimm, Differential role of Jak-STAT signaling in retinal degenerations. Faseb j 20, 2411–2413 (2006).

110. J. M. Mayrhofer, F. Haiss, D. Haenni, S. Weber, M. Zuend, M. J. P. Barrett, K. D. Ferrari, P. Maechler, A. S. Saab, J. L. Stobart, M. T. Wyss, H. Johannssen, H. Osswald, L. M. Palmer, V. Revol, C.-D. Schuh, C. Urban, A. Hall, M. E. Larkum, E. Rutz-Innerhofer, H. U. Zeilhofer, U. Ziegler, B. Weber, Design and performance of an ultra-flexible two-photon microscope for in vivo research. Biomed. Opt. Express 6, 4228–4237 (2015).

111. T. A. Pologruto, B. L. Sabatini, K. Svoboda, ScanImage: Flexible software for operating laser scanning microscopes. BioMedical Engineering OnLine 2, 13 (2003).

112. F. V. Moros, D. Amiet, R. M. Meister, A. von Faber-Castell, M. Wyss, A. S. Saab, P. Zbinden, B. Weber, L. Ravotto, A low-cost FPGA-based approach for pile-up corrected high-speed in vivo FLIM imaging. Neurophotonics 12, 025009 (2025).

113. S. C. Warren, A. Margineanu, D. Alibhai, D. J. Kelly, C. Talbot, Y. Alexandrov, I. Munro, M. Katan, C. Dunsby, P. M. French, Rapid global fitting of large fluorescence lifetime imaging microscopy datasets. PLoS One 8, e70687 (2013).

114. A. I. Erofeev, E. K. Vinokurov, O. L. Vlasova, I. B. Bezprozvanny, GCaMP, a Family of Single-Fluorophore Genetically Encoded Calcium Indicators. Journal of Evolutionary Biochemistry and Physiology 59, 1195–1214 (2023).

