## Supplemental material for "Photoreceptors have a dual dependency on both aerobic glycolysis and OXPHOS and diverge metabolically from other retinal neurons"

#### Figure S1

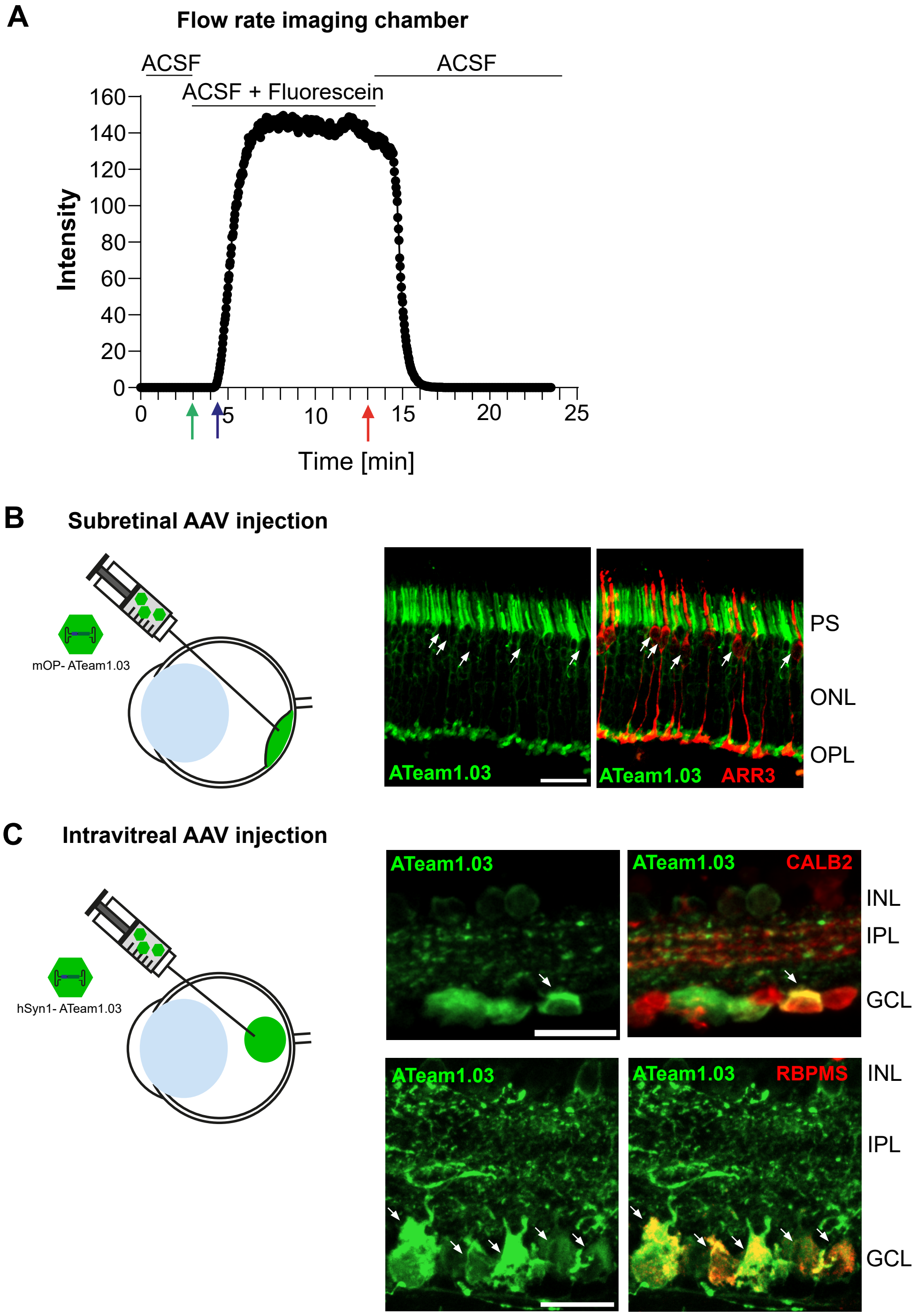

A
Flow rate imaging chamber
ACSF
ACSF + Fluorescein
ACSF
160
140
120
100
80
60
40
20
0
Intensity
0
5
10
15
20
25
Time [min]
B
Subretinal AAV injection
mOP- ATeam1.03
ATeam1.03
ATeam1.03 ARR3
PS
ONL
OPL
C
Intravitreal AAV injection
hSyn1- ATeam1.03
ATeam1.03
ATeam1.03 CALB2
INL
IPL
GCL
ATeam1.03
ATeam1.03 RBPMS
INL
IPL
GCL

#### Figure S2

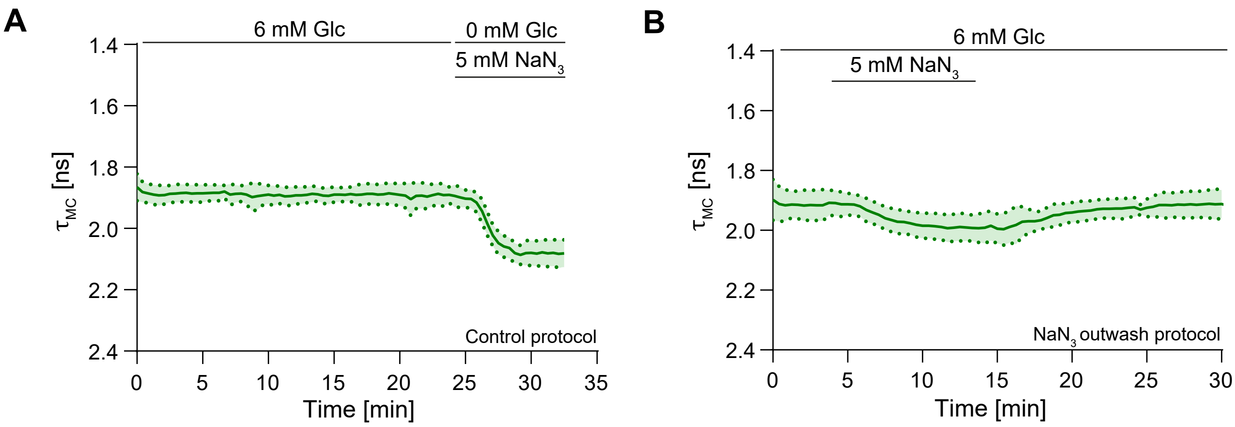

A
6 mM Glc
0 mM Glc
5 mM NaN3
1.4
1.6
1.8
2.0
2.2
2.4
τMC [ns]
Control protocol
0
5
10
15
20
25
30
35
Time [min]
B
6 mM Glc
5 mM NaN3
1.4
1.6
1.8
2.0
2.2
2.4
τMC [ns]
NaN3 outwash protocol
0
5
10
15
20
25
30
Time [min]

#### Figure S3

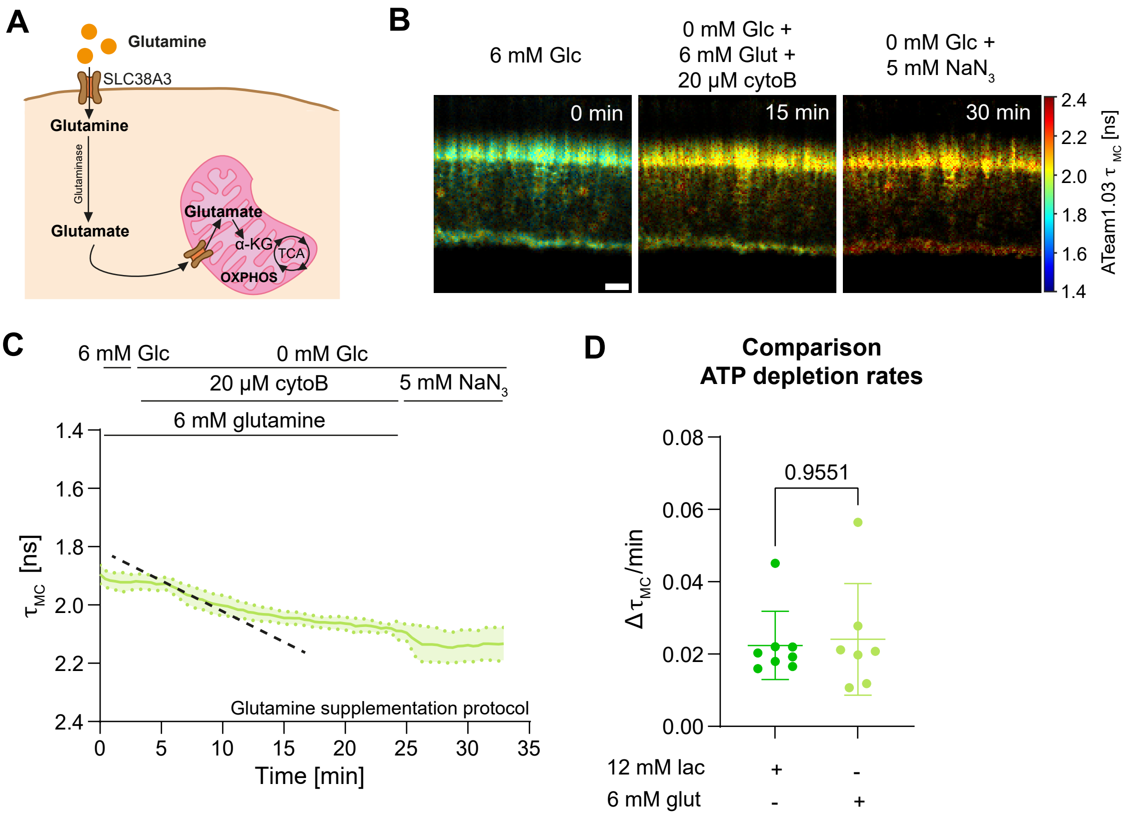
A
Glutamine
SLC38A3
Glutamine
Glutaminase
Glutamate
Glutamate
α-KG
TCA
OXPHOS
B
6 mM Glc
0 mM Glc +
6 mM Glut +
20 µM cytoB
0 mM Glc +
5 mM NaN3
0 min
15 min
30 min
2.4
2.2
2.0
1.8
1.6
1.4
ATeam1.03 τ MC [ns]
C
6 mM Glc
0 mM Glc
20 µM cytoB
5 mM NaN3
6 mM glutamine
τMC [ns]
1.4
1.6
1.8
2.0
2.2
2.4
Glutamine supplementation protocol
0
5
10
15
20
25
30
35
Time [min]
D
Comparison
ATP depletion rates
0.08
0.06
0.04
0.02
0.00
0.9551
Δ τMC/min
12 mM lac
+
-
6 mM glut
-
+

### Figure S4

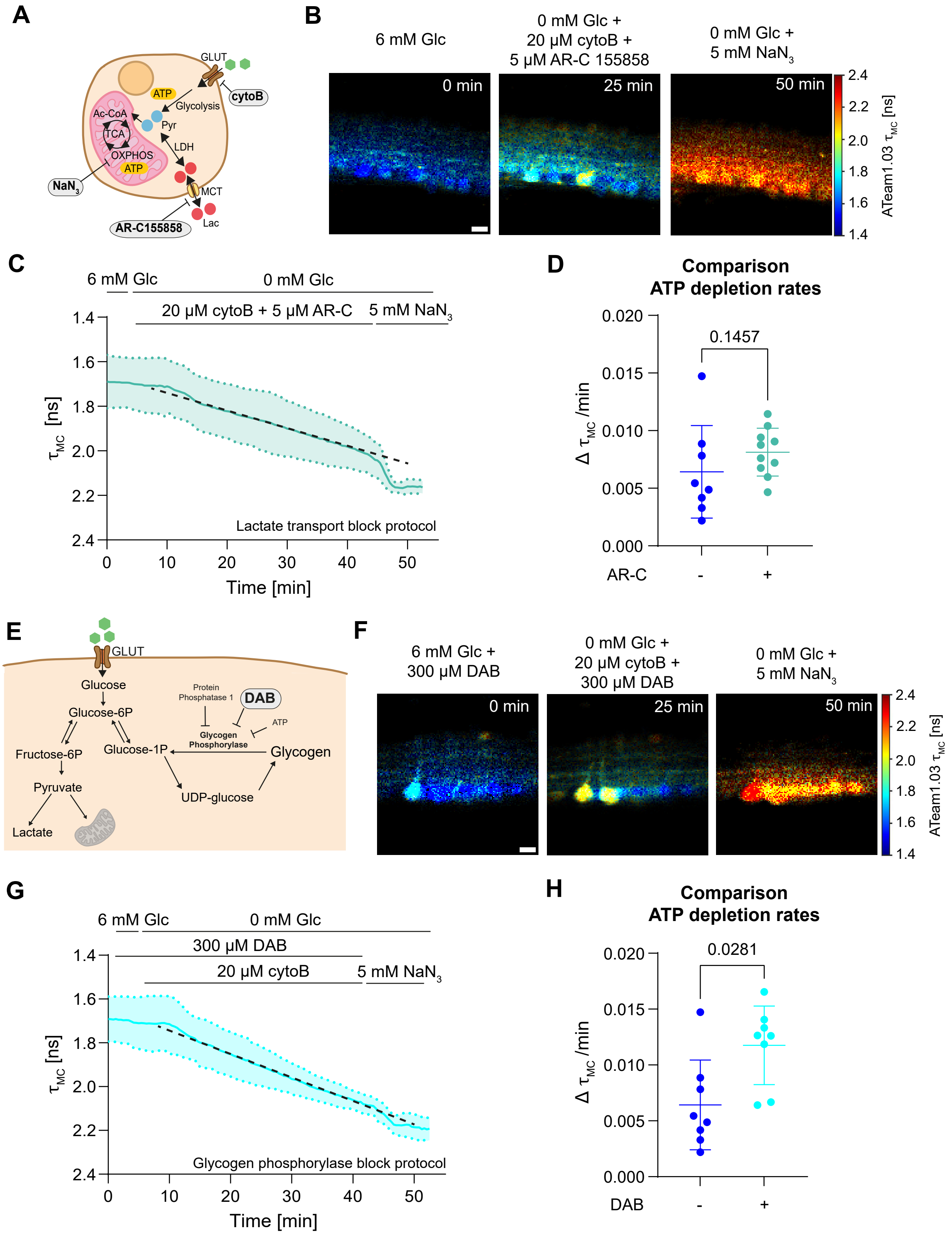

A
GLUT
cytoB
ATP
Glycolysis
Ac-CoA
TCA
Pyr
LDH
OXPHOS
ATP
NaN3
MCT
AR-C155858
Lac
B
6 mM Glc
0 mM Glc + 20 µM cytoB + 5 µM AR-C 155858
0 mM Glc + 5 mM NaN3
0 min
25 min
50 min
ATeam1.03 τMC [ns]
C
6 mM Glc
0 mM Glc
20 µM cytoB + 5 µM AR-C
5 mM NaN3
τMC [ns]
Lactate transport block protocol
Time [min]
D
Comparison ATP depletion rates
0.1457
Δ τMC /min
AR-C
-
+
E
GLUT
Glucose
Glucose-6P
Fructose-6P
Pyruvate
Lactate
Glucose-1P
Protein Phosphatase 1
DAB
ATP
Glycogen Phosphorylase
Glycogen
UDP-glucose
F
6 mM Glc + 300 µM DAB
0 mM Glc + 20 µM cytoB + 300 µM DAB
0 mM Glc + 5 mM NaN3
0 min
25 min
50 min
ATeam1.03 τMC [ns]
G
6 mM Glc
0 mM Glc
300 µM DAB
20 µM cytoB
5 mM NaN3
τMC [ns]
Glycogen phosphorylase block protocol
Time [min]
H
Comparison ATP depletion rates
0.0281
Δ τMC /min
DAB
-
+

#### Figure S5

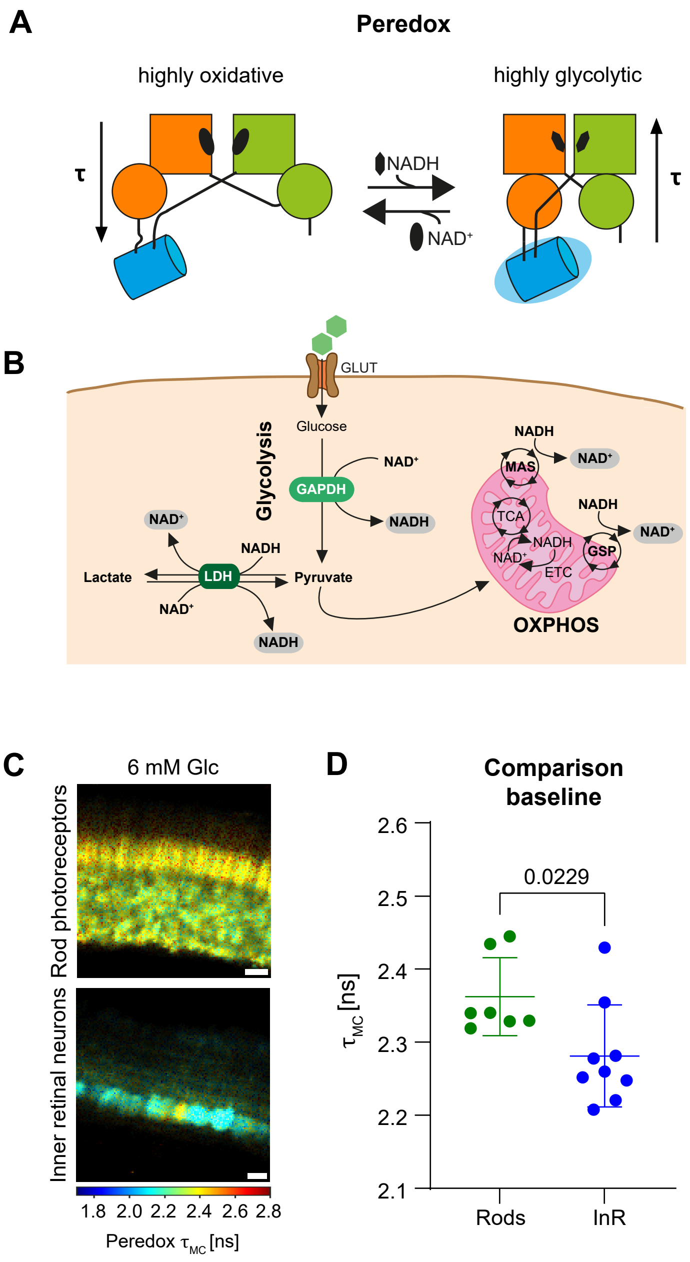
A
Peredox
highly oxidative
highly glycolytic
τ
NADH
NAD+
τ
B
GLUT
Glucose
Glycolysis
NAD+
GAPDH
NADH
NAD+
NADH
Lactate
LDH
Pyruvate
NAD+
NADH
NADH
NAD+
MAS
TCA
NADH
NAD+
NADH
NAD+
GSP
ETC
OXPHOS
C
6 mM Glc
Rod photoreceptors
Inner retinal neurons
1.8 2.0 2.2 2.4 2.6 2.8
Peredox τMC [ns]
D
Comparison baseline
0.0229
τMC [ns]
2.6
2.5
2.4
2.3
2.2
2.1
Rods
InR

#### Figure S6

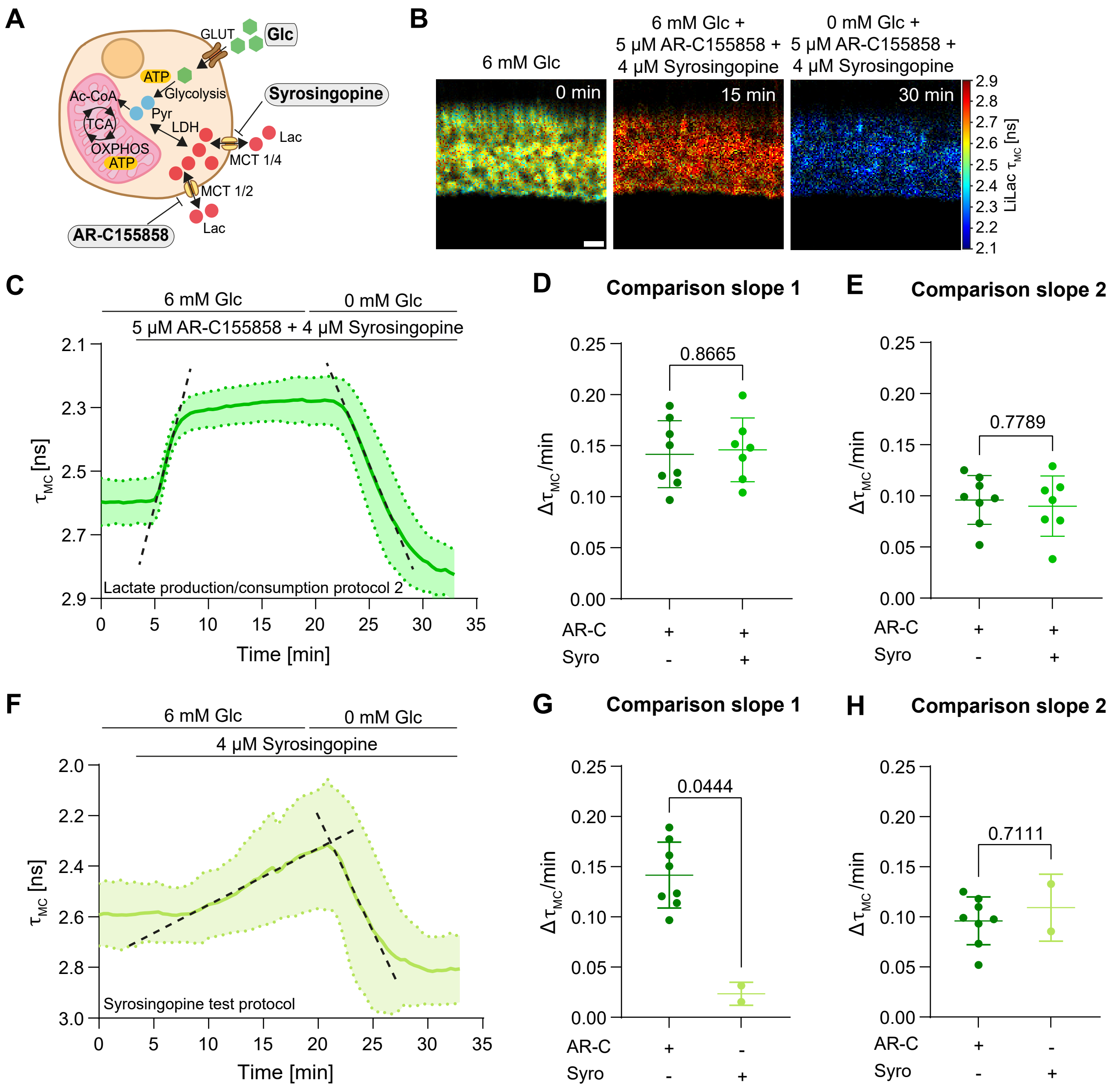

A
GLUT
Glc
ATP
Glycolysis
Syrosingopine
Ac-CoA
TCA
Pyr
LDH
Lac
OXPHOS
ATP
MCT 1/4
MCT 1/2
AR-C155858
Lac
B
6 mM Glc
6 mM Glc + 5 μM AR-C155858 + 4 μM Syrosingopine
0 mM Glc + 5 μM AR-C155858 + 4 μM Syrosingopine
0 min
15 min
30 min
LiLac $\tau_{MC}$ [ns]
C
6 mM Glc
0 mM Glc
5 μM AR-C155858 + 4 μM Syrosingopine
$\tau_{MC}$ [ns]
Lactate production/consumption protocol 2
Time [min]
D
Comparison slope 1
0.8665
$\Delta\tau_{MC}$/min
AR-C
Syro
E
Comparison slope 2
0.7789
F
6 mM Glc
0 mM Glc
4 μM Syrosingopine
Syrosingopine test protocol
G
Comparison slope 1
0.0444
H
Comparison slope 2
0.7111

#### Figure S7

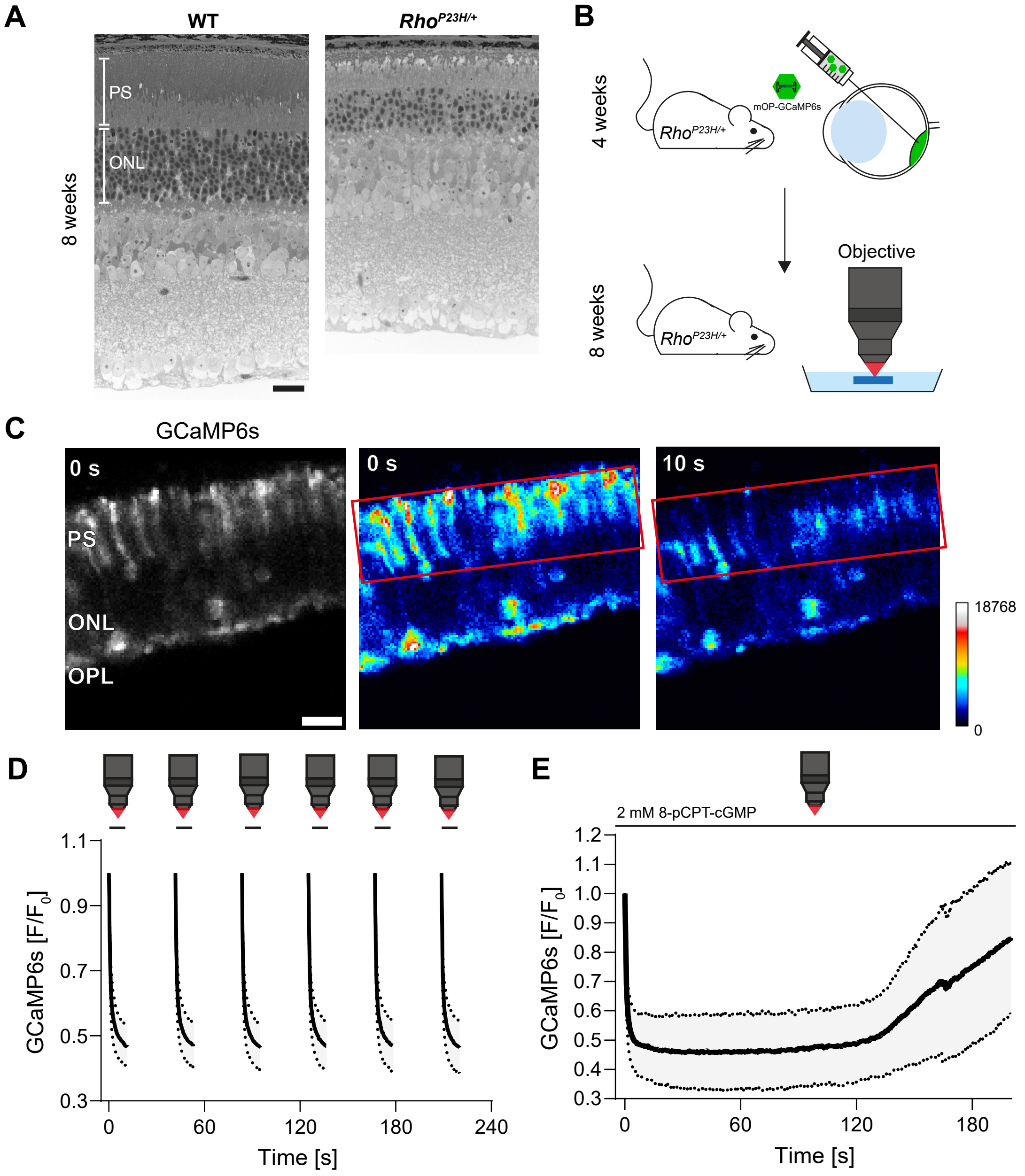
A
WT
Rho P23H/+
PS
ONL
8 weeks
B
4 weeks
Rho P23H/+
mOP-GCaMP6s
8 weeks
Objective
C
GCaMP6s
0 s
PS
ONL
OPL
0 s
10 s
18768
0
D
1.1
0.9
0.7
0.5
0.3
GCaMP6s [F/F0]
0
60
120
180
240
Time [s]
E
2 mM 8-pCPT-cGMP
1.2
1.1
1.0
0.9
0.8
0.7
0.6
0.5
0.4
0.3
GCaMP6s [F/F0]
0
60
120
180
Time [s]

**Table S1:** Drugs used for 2P or 2P-FLIM experiments

| Drug | Stock conc. | Solvent | Final conc. | Source, cat. no. |
|---|---|---|---|---|
| Sodium azide ($NaN_3$) | 5 M | $H_2O$ | 5 mM | Sigma-Aldrich, S2002 |
| 8-pCPT-cGMP; 8- (4-Chlorophenylthio)guanosine- 3’, 5’- cyclic monophosphate | 20 mg | ACSF | 2 mM | Enzo BioLog (Farmingdale, NY, USA), C009-50 |
| DAB; 1,4-Dideoxy-1,4-imino-D-arabinitol hydrochloride | 100 mM | $H_2O$ | 300 µM | Santa Cruz (Dallas, TX, USA), sc-220553A |
| AR-C155858 | 10 mM | DMSO | 5 µM | BioTechne Torcis (Bristol, UK), 4960/1 |
| cytoB; Cytochalasin B | 10 mM | DMSO | 20 µM | BioTechne Torcis, 5474/10 |
| Syrosingopine | 10 mM | DMSO | 4 µM | Merck Millipore Chemicals, SML1908 |
